# A novel method for an unbiased estimate of cross-ancestry genetic correlation using individual-level data

**DOI:** 10.1101/2021.09.16.460619

**Authors:** Md. Moksedul Momin, Jisu Shin, Soohyun Lee, Buu Truong, Beben Benyamin, S. Hong Lee

## Abstract

Cross-ancestry genetic correlation is an important parameter to understand the genetic relationship between two ancestry groups for a complex trait. However, existing methods cannot properly account for ancestry-specific genetic architecture, which is diverse across ancestries, producing biased estimates of cross-ancestry genetic correlation. Here, we present a method to construct a genomic relationship matrix (GRM) that can correctly account for the relationship between ancestry-specific allele frequencies and ancestry-specific causal effects. Through comprehensive simulations, we show that the proposed method outperforms existing methods in the estimations of SNP-based heritability and cross-ancestry genetic correlation. The proposed method is further applied to six anthropometric traits from the UK Biobank data across 5 ancestry groups. One of our findings is that for obesity, the estimated genetic correlation between African and European ancestry cohorts is significantly different from unity, suggesting that obesity is genetically heterogenous between these two ancestry groups.

## Introduction

Complex traits are polygenic and influenced by environmental factors^1,2^, which can be distinguished from Mendelian traits that are regulated by single or few major genes and minimal environmental influences. In humans, causal loci and their effects on complex traits are dynamically distributed across populations such that the same trait can be genetically heterogenous between two ancestry groups. Studies of cross-ancestry genetic correlation are therefore important to understand the genetic relationship between two ancestry groups when investigating a complex trait^2^. Understanding genetic correlation between two ancestry groups can inform us on how well we can predict a complex disease in one ancestry group based on the information on another.

Genome wide association studies (GWAS) have been successful in identifying variants associated with causal loci, and can also provide a useful resource to investigate cross-ancestry genetic correlations. Using GWAS datasets, genomic relationships between samples across multiple ancestry groups can be constructed based on genome-wide single nucleotide polymorphisms (SNPs), which can be fitted with phenotypes across ancestries in a statistical model^3,4^. This paradigm-shift approach to estimate cross-ancestry genetic correlations have greatly increased the potential to dissect the latent genetic relationship between ancestry groups for complex traits.

Most GWAS have focused on European ancestry samples (> 80%)^5–7^ although Europeans represent only 16% of the global population^8–10^. Because of large samples, the estimated SNP associations in Europeans are far more accurate than those in other ancestries. As a matter of fact, the performance of polygenic risk prediction depends on the accuracy of estimated SNP associations, causing a disparity in genetic prediction across populations^5^. Therefore, cross-ancestry genetic studies are urgently required to bridge the disparity, e.g. estimated cross-ancestry genetic correlations may be able to inform if SNP effects estimated in European ancestry samples can be useful to predict the phenotypes of samples in other ancestry groups.

To date, several studies have been undertaken to estimate cross-ancestry genetic correlations between diverse ancestry groups for a range of complex traits^2,3,11–13^. For example, Yang et al.^4^ estimated the cross-ancestry genetic correlation between East Asians and Europeans for ADHD, where ancestry-specific allele frequencies were used to standardise samples’ genotype coefficients in estimating their genomic relationships. This method of cross-ancestry genomic relationships has been widely used for cross-ancestry genetic studies^3,14^. However, the method cannot account for the trait genetic architecture specific to each ancestry group, which is a function of the relationship between the genetic variance and allele frequency (one important aspect of a heritability model^15,16^). With an incorrect heritability model, estimated genetic variances within and covariance between ancestry groups are biased^15,16^, and hence the cross-ancestry genetic correlations cannot be correctly estimated. Another method based on GWAS summary statistics has been introduced (Popcorn)^2^. However, this method also cannot correctly account for diverse genetic architectures across ancestry groups.

Here, we develop a novel method that can properly account for ancestry-specific genetic architecture and ancestry-specific allele frequency in estimating a genomic relationship matrix (GRM). In addition, we revisit the existing theory to correct mean bias in genomic relationships. In simulations, the SNP-based heritability and cross-ancestry genetic correlation estimated from our proposed method are shown to be unbiased, whereas other existing methods generate biased estimates. We apply the proposed method to six anthropometric traits from the UK Biobank data that include standing height, body mass index (BMI), waist circumference, hip circumference, waist-hip ratio, and weight. For each trait, we estimate SNP-based heritabilities and cross-ancestry genetic correlations across 5 ancestry groups, i.e., white British, other Europeans, Asian, African, and mixed ancestry cohorts.

## Results

### Overview

We used publicly available data from the UK Biobank. Participants of the UK Biobank were stratified into multiple ancestries (white British, other European, Asian, African, and mixed ancestry cohorts) according to their underlying genetic ancestry based on a principal component analysis (see **Supplementary Figure S1**)^17^. In each ancestry group, we assume that the relationship between genetic variance and allele frequencies (heritability model) varies, i.e. the genetic variance at the *i^th^* genetic variant can be expressed as:

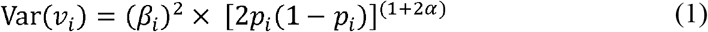

where *β_i_* and *p_i_* are the causal effects and the reference allele frequency of the *i^th^* genetic variant, *v_i_*, and *α* is the scale factor determining the genetic architecture of complex traits in each ancestry groups. Note that with *α* = 0, equation (1) becomes the classical model of Falconer and Mackay (1996)^18^. By assuming that the genetic variance of causal variant is constant across minor allele frequencies (MAF) spectrum, a heritability model with *α* = −0.5 has been widely used^19–21^. However, Speed et al.^15,16,22^ reported a different *α* value, e.g., *α* = −0.125 for anthropometric traits, using multiple European cohorts. While it is intuitive that *α* values can be varied across ancestry groups, it has not been well studied.

First, we determine an optimal *α* value for each ancestry group, comparing model fits (maximum log-likelihood) of various heritability models with different *α* values for the 6 anthropometric traits from the UK Biobank (see Methods). Second, we simulate phenotypes based on the UK Biobank genotypic data to assess if SNP-based heritability and cross-ancestry genetic correlation are unbiasedly estimated. In the simulation, various *α* values are used to generate causal effects of SNPs in various ancestry groups, and the correlation of SNP effects between ancestry groups varies between 0 and 1 (see Methods). For simulated phenotypes of multiple ancestry groups, we estimate SNP-based heritability and cross-ancestry genetic correlation, using bivariate GREML^20,21,23^ with four existing methods to construct GRM as shown in **Table 1**. In addition to the existing methods, we use a novel method to estimate GRM in the cross-ancestry analysis in which we use ancestry-specific *α* value and ancestry-specific allele frequency, so that the estimation model matches with the ancestry-specific genetic architecture (see Methods). The equation for the proposed method can be written as

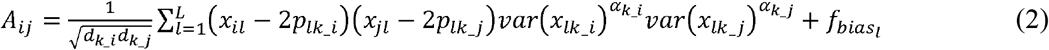

where *x_il_* and *x_jl_* are SNP genotypes for the *i^th^* and *j^th^* individuals at the *l*^th^ SNP, *p_lk_i_* and *p_lk_j_* are the allele frequencies at the *l*^th^ SNP (*l* = 1 – L, where L is the number of SNPs) estimated in the two ancestry groups, *k_i* and *k_j*, to which the *i^th^* and *j^th^* individuals belongs, and *α_k_i_* and *α_k_j_* are the scale factors for the two ancestry groups, *x_lk_i_* and *x_lk_j_* are all individual genotypes at the *l*^th^ SNP in the two ancestry groups, *d_k_i_* is the expectation of the diagonals, 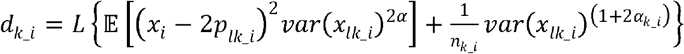, and *f_bias_l__* is the bias factor at the *l*th SNP, 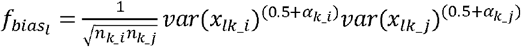 where *n_k_i_* and *n_k_j_* are the number of individuals in the two ancestry groups. The term, *f_bias_l__*, can correct for the mean bias in the existing equations^19,20^ (see Methods and **Supplementary Table 1**). In addition, we note that using var(*x_l_*), instead of its expectation (2*p_l_*(1 – *p_l_*)), is more robust^24^ (**Supplementary Table 2**).

**Table 1.** Four existing methods to estimate cross-ancestry genetic correlation, compared to the proposed method.

| Methods | Scale factor<br>($\alpha$) | SNPs | Allele frequency | Reference | Equation<br>No. |
| --- | --- | --- | --- | --- | --- |
| GRM1 | -0.5 | All<sup>a</sup> | overall-average<sup>b</sup> | 3,19,20 | (5, 7) |
| GRM2 | -0.5 | common only<sup>c</sup> | overall average | 3,19,20 | (5,7) |
| GRM3 | -0.5 | All | ancestry-specific<sup>d</sup> | 3,4 | (10) |
| GRM4 | -0.5 | common only | ancestry-specific | 3,4 | (10) |
| Proposed<br>method | ancestry-specific<sup>e</sup> | common only | ancestry-specific |  | (2) |

<sup>a</sup>Using all SNPs from both ancestry groups. <sup>b</sup>Allele frequencies estimated from the combined population of both ancestry groups when scaling the genotypes. <sup>c</sup>Using only common SNPs presented for both ancestry groups after QC including ancestry specific MAF > 0.01. <sup>d</sup>Allele frequencies estimated from each population when scaling the genotypes. <sup>e</sup>Different $\alpha$ value specific to each ancestry group can be used when scaling the genotypes.

Finally, we analyse real data using the proposed method (equation 2) to estimate SNP-based heritability and cross-ancestry genetic correlation for 6 anthropometric traits across different ancestries using bivariate GREML.

### Determination of scale factor (α) across ancestries

We compared the Akaike information criteria (AIC) values of heritability models with varying *α* values to determine which *α* value provides the best model fit (see Methods), which is analogue to the approach of Speed et al.^16^. In terms of LD weights, we contrasted two kinds of heritability models, i.e. GCTA-*α* vs. LDAK-thin-*α* model. GCTA-*α* model has no LD weights, whereas LDAK-thin-*α* model explicitly considers LD among SNPs.

When using GCTA-*α* model, we observed that AIC values with *α* = −0.25, −0.125, −0.625, −0.75 and −0.825 were lowest for white British, other European, Asian, African, and mixed ancestry cohorts, respectively (**Figure 1 and Supplementary Tables 3–7**). When considering LDAK-thin-*α* model, we estimated optimal *α* values as −0.25, −0.125, −0.50, −0.625 and −0.75 for white British, other European, Asian, African, and mixed ancestry cohorts, respectively (**Supplementary Figure 2 and Supplementary Tables 3–7**).

**Figure 1.**
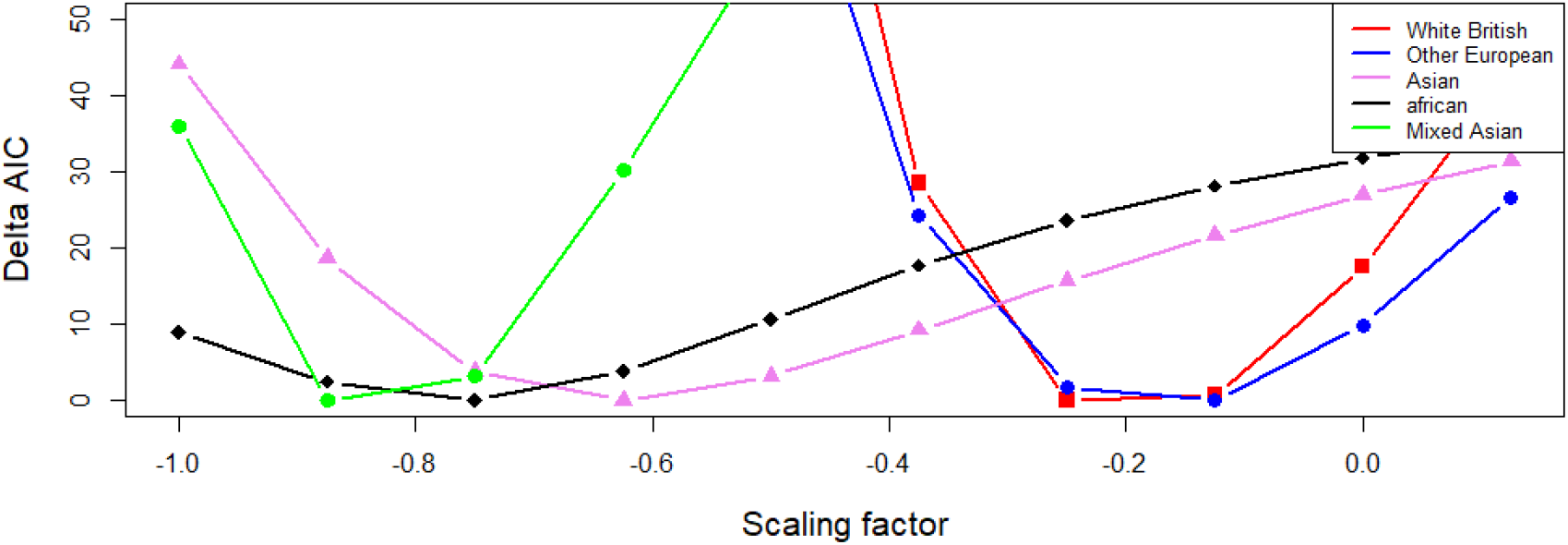
Determining optimal scale factors (*α*) for 5 different ancestry groups using GCTA-*α* model. GCTA-*α* model assumes that all SNPs have an equal contribution to the heritability estimation and *α* value varies across ancestries^15,16^. ΔAIC values from GCTA-*α* models are plotted against scaling factors, *α*, for each ancestry group. The lowest AIC (i.e. ΔAIC=0) indicates the best model. The sample sizes are 30,000, 26,457, 6,199, 6,179 and 11,797 for white British, other European, Asian, African, and mixed ancestry groups, respectively.

When comparing GCTA-*α* and LDAK-thin-*α* models, the AIC value of GCTA-*α* model was much smaller than that of LDAK-thin-*α* model for white British or other European ancestry cohort (**Supplementary Table 8**). For Asian ancestry cohort, the AIC of GCTA-*α* model was slightly lower than LDAK-thin-*α* model when using the best *α* value = −0.625. In contrast, the AIC of LDAK-thin-*α* model was generally lower than that of GCTA-*α* model for African or mixed ancestry cohort.

### Method validation by simulation

We simulated phenotypes based on the real genotypic data of multiple ancestry groups where the estimated *α* value for each ancestry group was used to generate SNP effects that were correlated between ancestry groups (Methods). In this simulation, we do not consider associations between the causal effects and LD structure for any SNP, i.e. LDAK simulation model, because LDAK model was not particularly plausible for the genetic architecture of the traits especially for white British, other European and Asian ancestry cohorts (**Supplementary Table 8**) and LDAK simulation model was not feasible for a bivariate framework. When using simulation models with *α* = −0.5 for all ancestry groups, estimated SNP-based heritabilities were mostly unbiased for all the methods, GRM1-4 (**Supplementary Table 9**). Estimated genetic correlations from GRM3 and 4 were unbiased for all scenarios (**Figure 2 and Supplementary Table 9**). However, estimated genetic correlations from GRM1 and 2 were biased when the true genetic correlation was high (**Figure 2 and Supplementary Table 9**). This shows that estimated cross-ancestry genetic correlation can be biased unless ancestry-specific allele frequencies were properly accounted for.

**Figure 2.**
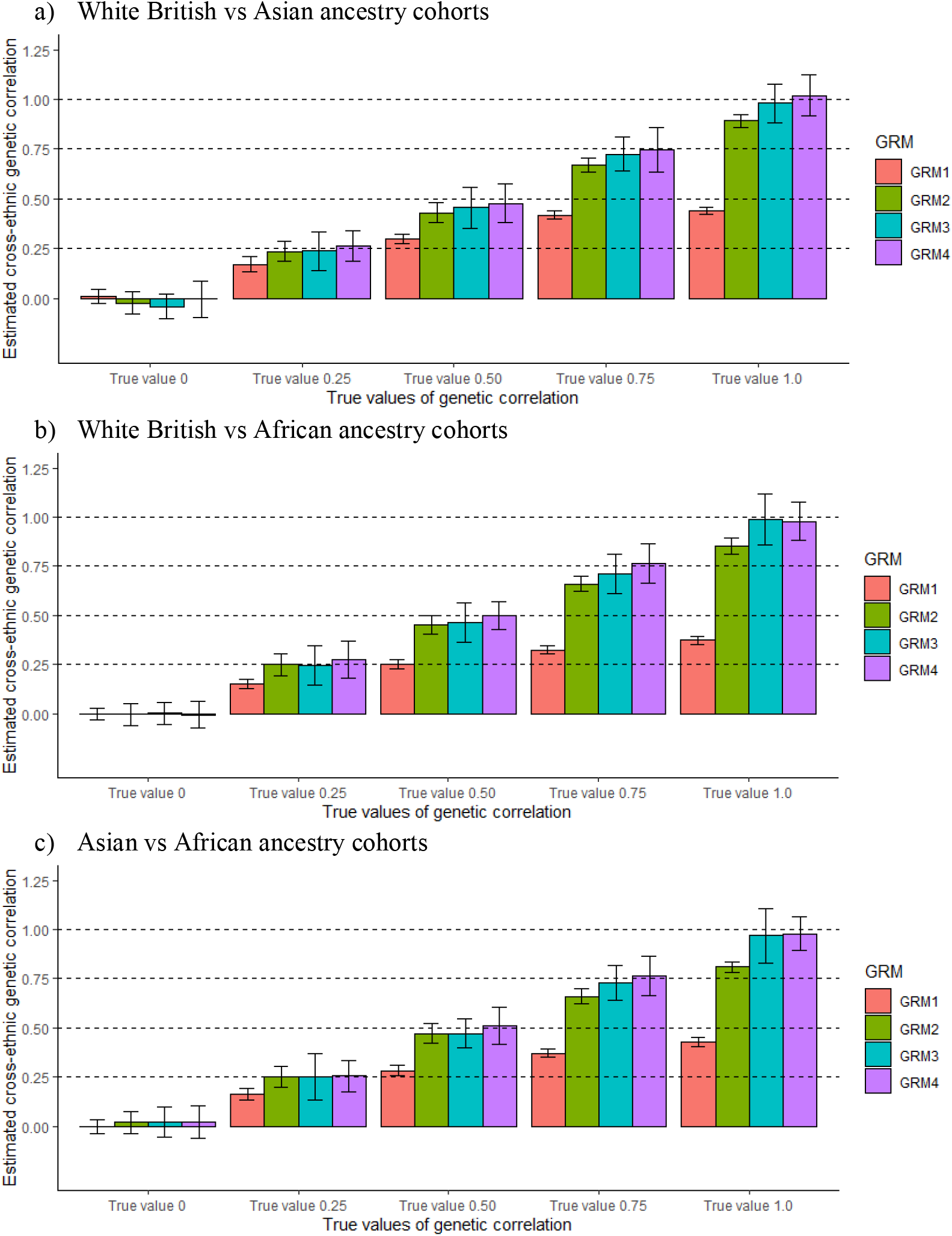
Estimated cross-ancestry genetic correlations from four existing methods when *α* is fixed to −0.5 for both simulation and estimation. Height of each bar is the estimated cross-ancestry genetic correlation and error bar indicates 95% confidence interval (CI) for 500 replicates. In our simulations, we used three combinations for estimating cross-ancestry genetic correlation (white British vs. Asian, white British vs. African and Asian vs. African ancestry cohorts). In each ancestry group, we used 500,000 SNPs that were randomly selected from HapMap3 SNPs after QC. To simulate phenotypes, we selected a random set of 1,000 SNPs as causal variants, which were presented for both ancestry groups. We used *α* = −0.5 when scaling the causal effects by ancestry-specific allele frequency in each ancestry group. Various values of genetic correlation were considered (0, 0.25, 0.50, 0.75 and 1.0). In the estimation, the four methods (GRM1 – 4) used *α* = −0.5 (standard scale factor in GRM estimation). GRM1 and 3 used all available SNPs from both ancestry groups (791581, 812332 and 777894 for figure panels a, b and c) whereas GRM2 and 4 used only the set of SNPs common between two ancestry groups (208419, 187668 and 222106 for a, b, and c). When scaled with *α* = −0.5, GRM1 and 2 used allele frequency averaged between two ancestry groups whereas GRM3 and 4 used ancestry-specific allele frequency estimated from each ancestry group (see **Table 1**).

To mimic real data, we simulated phenotypes based on the real genotypes of multiple ancestry groups, using realistic *α* values (*α* = −0.25 for white British ancestry cohorts, *α* = −0.625 for Asian ancestry cohorts and *α* = −0.75 for African ancestry cohorts) instead of using a constant *α* value across ancestries (Methods). For these simulated phenotypes, estimated cross-ancestry genetic correlations using four existing methods were biased when the true genetic correlation was 0.5 or higher between white British and African ancestry cohorts or between Asian and African ancestry cohorts. Biased estimates were still observed even when ancestry-specific allele frequencies were considered in GRM3 and 4 (**Figure 3b and 3c**). As expected, estimated SNP-based heritability were mostly biased because of mis-specified *α* values when estimating GRMs (**Supplementary Figure 3 and Supplementary Table 10**).

**Figure 3.**
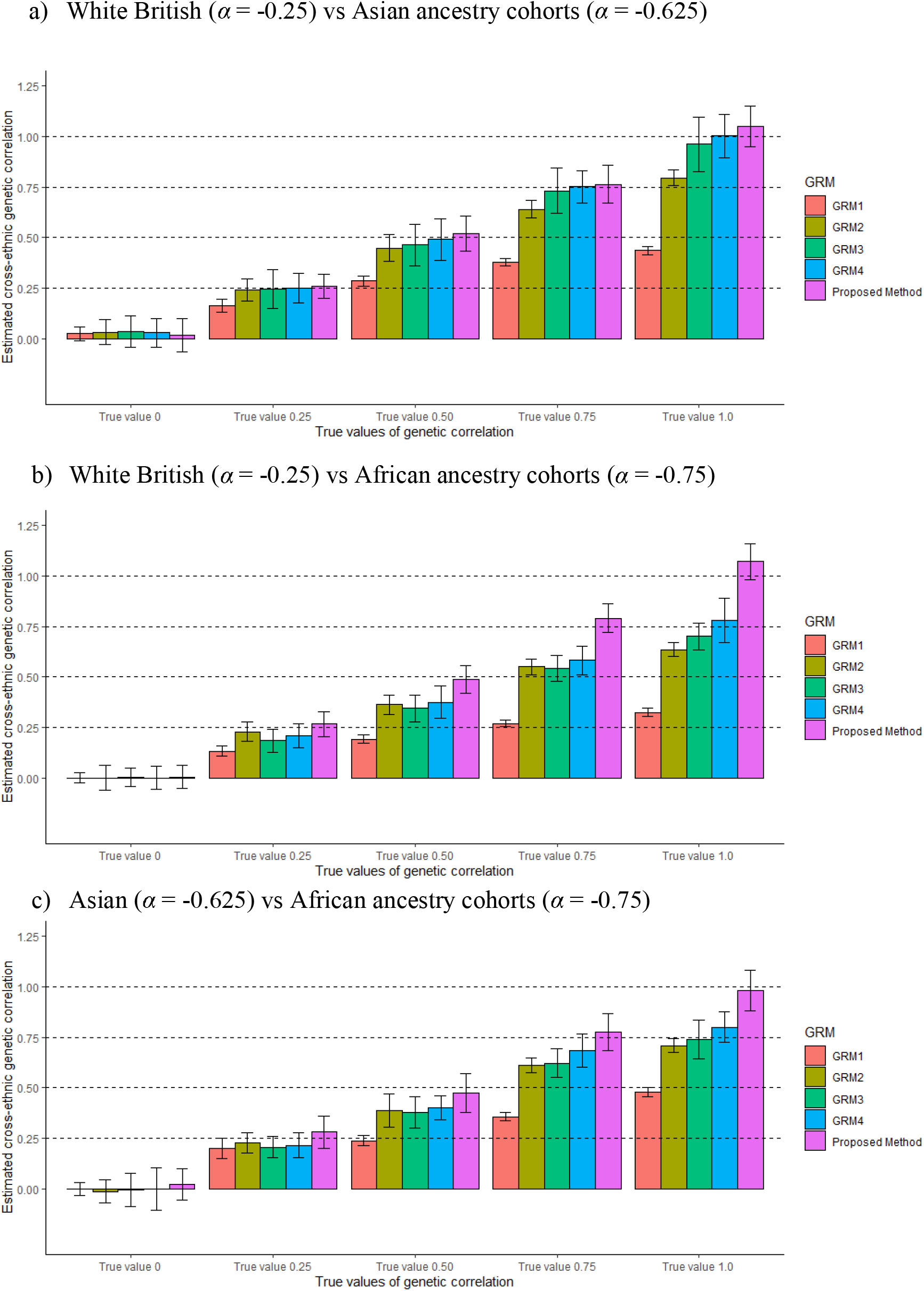
Estimated cross-ancestry genetic correlations from four existing methods when varying *α* values across ancestry groups. Height of each bar is the estimated cross-ancestry genetic correlation and error bar indicates 95% confidence interval (CI) for 500 replicates. In our simulation, we used three combinations for estimating cross-ancestry genetic correlation (white British vs. Asian, white British vs. African and Asian vs. African ancestry cohorts). In each ancestry group, we used 500,000 SNPs that were randomly selected from HapMap3 SNPs after QC. To simulate phenotypes, we selected a random set of 1,000 SNPs as causal variants, which were presented for both ancestry groups. We used various *α* values that were specific to ancestries (*α* = −0.25, −0.625 and −0.75 for white British, Asian and African ancestry cohorts, respectively) when scaling the causal effects by ancestry-specific allele frequency in each ancestry group. Various values of genetic correlation were considered (0, 0.25, 0.50, 0.75 and 1.0). In the estimation, we used existing methods (GRM1 – 4) that used the standard scale factor *α* = −0.5 in GRM estimation. GRM1 and 3 used all available SNPs from both ancestry groups (791581, 812332 and 777894 for figure panels a, b and c) whereas GRM2 and 4 used only the set of SNPs common between two ancestry groups (208419, 187668 and 222106 for a, b, and c). When scaled with *α* = −0.5, GRM1 and 2 used allele frequency averaged between two ancestry groups whereas GRM3 and 4 used ancestry-specific allele frequency estimated from each ancestry group. In addition, we applied the proposed method that used ancestry-specific *α* value and ancestry-specific allele frequency in GRM estimation.

For the same simulated phenotypes, we applied the proposed method that accounts for ancestry-specific allele frequency and ancestry-specific *α* values (equation 2) and found that it provided unbiased estimates for both SNP-based heritability and cross-ancestry genetic correlation (**Figure 3 and Supplementary Table 11)**. It was noted that the proposed method was robust to different numbers of causal SNPs (**Supplementary Table 12**).

Popcorn software, a GWAS summary statistics-based method for cross-ancestry genetic analysis, also generated biased estimates of cross-ancestry genetic correlation when using realistic *α* values in the simulation (**Supplementary Figure 4**). This agrees with Brown et al.^2^, that popcorn estimates can be deviated from the true values when using alternative genetic architectures (or heritability models).

### SNP-based Heritability (h^2^) estimates for anthropometric traits using real data

The estimated SNP-based heritability for each of the anthropometric traits was presented in **Figure 4**. The estimated SNP-based heritability of standing height was found to be highest in white British ancestry cohort (0.502, SE= 0.012) and lowest in African ancestry cohort (0.246, SE= 0.053). Heritability estimation in African ancestry cohort was significantly lower than white British ancestry cohorts (*p*-value = 2.47e-06), other European ancestry cohorts (*p*-value = 2.30e-05) and Asian ancestry cohorts (*p*-value = 1.14e-03). The estimates for BMI and waist and hip circumference were generally high in African ancestry cohorts and low in other European ancestry cohorts (**Figure 4**). The estimated SNP-based heritability ranged from 0.231 (other European ancestry cohorts) to 0.322 (Asian ancestry cohorts) for weight, and from 0.043 (African ancestry cohorts) to 0.153 (Asian ancestry cohorts) for waist-hip ratio. In contrast to height, there was no significant difference among ancestry-specific heritability estimates for BMI, waist circumference, hip circumference, waist hip ratio or weight.

**Figure 4.**
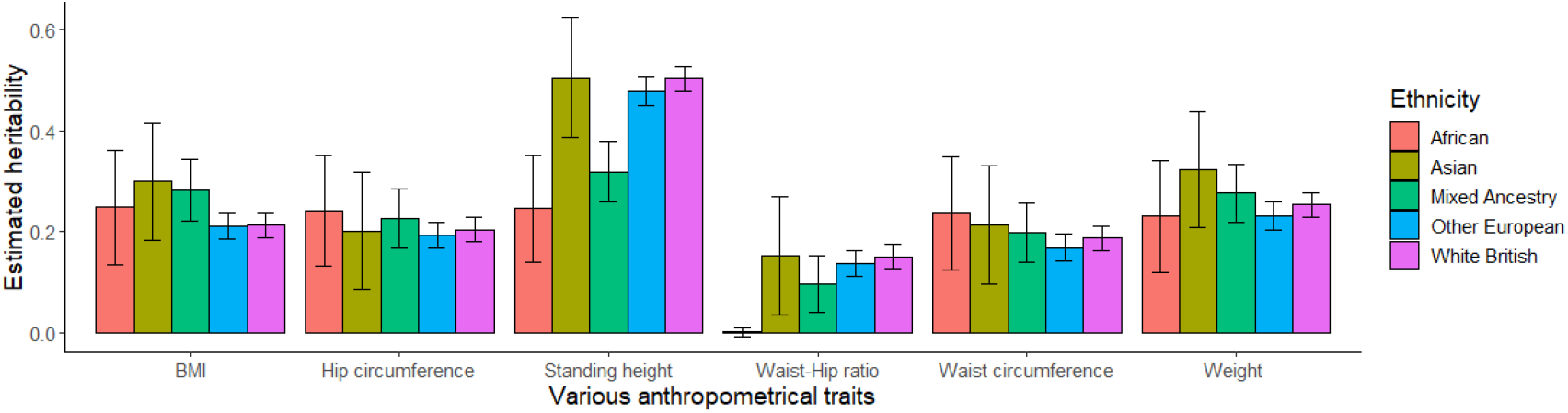
Estimated SNP-based heritability for different anthropometric traits across ancestries. The main bars indicate estimated SNP-based heritabilities and the error bars indicate 95% confidence intervals.

### Cross-ancestry genetic correlations for anthropometric traits

Estimated cross-ancestry genetic correlations (*r_g_*) between ancestry groups for 6 anthropometric traits are shown in **Figure 5**. For BMI, the estimated genetic correlations between African and white British ancestry cohorts (*r_g_* = 0.672; SE =0.131; *p*-value = 1.22e-02) and between African and other European ancestry cohorts (*r_g_* = 0.549; SE= 0.134; *p*-value= 7.63e-04) were significantly different from 1 (**Figure 5a and Supplementary Table 13**). This indicated that BMI is a genetically heterogenous between African and European ancestry cohorts. Estimated genetic correlations between African and Asian or European and Asian ancestry cohorts were low, but not significantly different from 1 (i.e. no evidence of genetic heterogeneity). As expected, the estimated genetic correlation between white British and other European was not significantly different from 1 (*r_g_* = 1.081, SE = 0.043, *p*-value = 5.96e-02).

**Figure 5.**
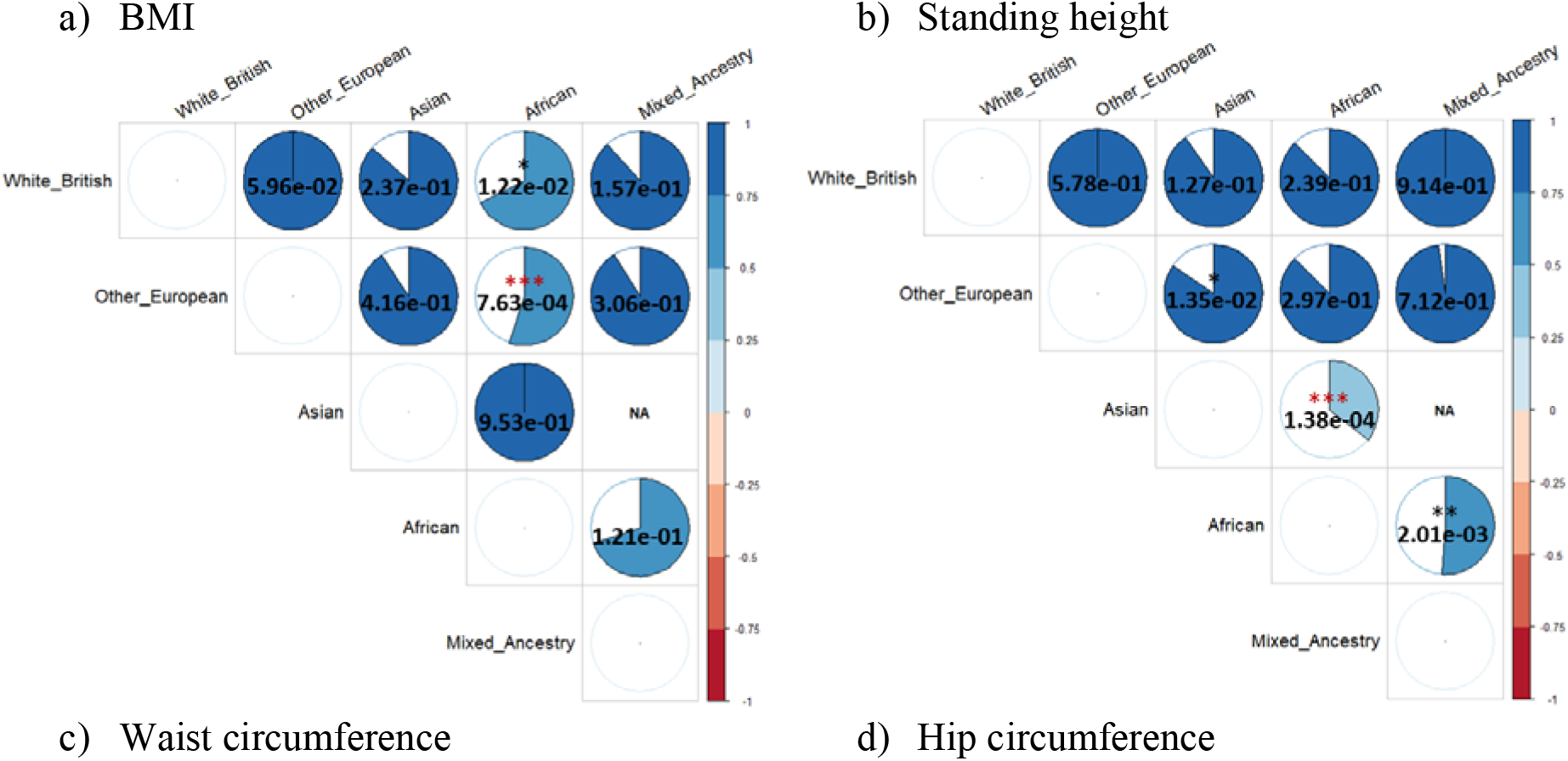
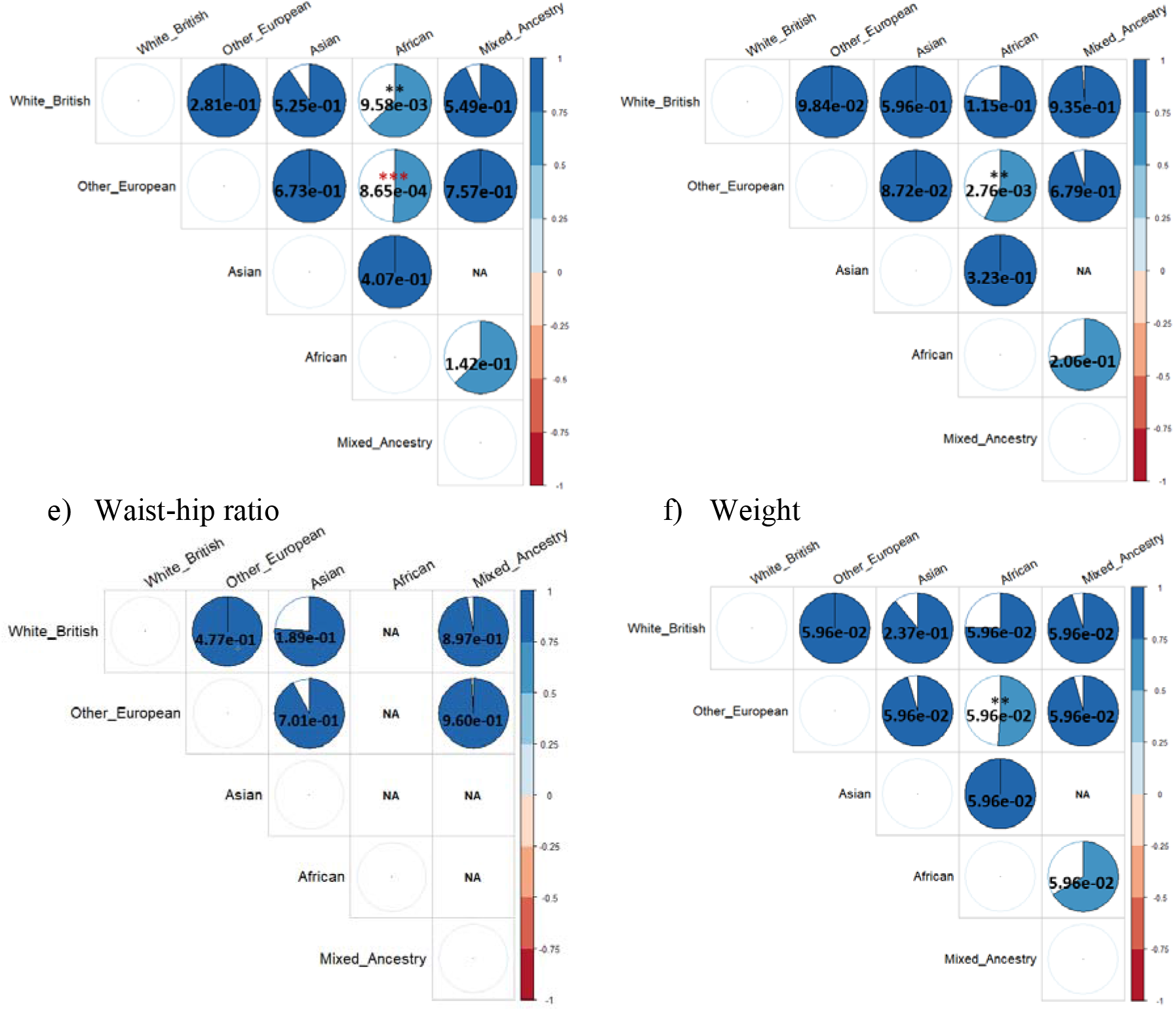
Estimated cross-ancestry genetic correlations for anthropometric traits. The colour and size of each pie chart indicates the magnitude of estimated cross-ancestry genetic correlations. The value in each pie chart is a *p*-value (*, ** and *** indicates p value < 0.05, < 0.01 and <0.001, respectively) testing the null hypothesis of *r_g_*=1. Coloured asterisk indicates significantly different from 1 after Bonferroni correction (0.05/54). Non-estimable parameter is shown as NA, which was due to one ancestry group is nested within the other ancestry groups or estimated SNP-*h^2^* is zero for one ancestry group.

For height, we observed a significant genetic heterogeneity between Asian and African ancestry cohorts (*r_g_* = 0.356; SE= 0.69; p-value = 1.38e-04), between other European and Asian ancestry cohorts (*r_g_* = 0.847; SE= 0.062; *p*-value = 1.35e-02) and between African and mixed ancestry cohorts (*r_g_* = 0.512; SE= 0.158; *p*-value = 2.01e-03) (**Figure 5b and Supplementary Table 14**). Although estimated genetic correlations were lower than 1, there were no significant evidence for genetic heterogeneity between white British and Asian ancestry cohorts (*r_g_* = 0.904; SE= 0.063; *p*-value =1.27e-01), and between white British and African ancestry cohorts (*r_g_* = 0.876; SE= 0.118; *p*-value =2.93e-01). White British and other European ancestry cohorts were observed to be genetically homogeneous for the trait (*r_g_* = 1.01; SE= 0.018; *p*-value = 5.78e-01).

Estimated cross-ancestry genetic correlations for waist circumference and hip circumference showed a similar pattern with BMI, i.e., there was a significant evidence for genetic heterogeneity between white British and African ancestry cohorts and between other Europeans and African ancestry cohorts (**Figure 5c and 5d; Supplementary Tables 15 and 16**). The estimated genetic correlation between African and mixed ancestry cohorts was low, but not significantly different from 1. As the same as in BMI and height, there was no genetic heterogeneity between white British and other European ancestry cohorts for both waist circumference and hip circumference.

For weight, the estimated genetic correlation between African and other European ancestry cohorts was significantly different from 1 (*r_g_* =0.624; SE=0.139; *p*-value = 6.83e-03), indicating a significant genetic heterogeneity between these two ancestry groups (**Figure 5f and Supplementary Table 18**). Although the estimations are not significant, we have estimated lower genetic correlations (far from 1) between white British and African ancestry cohorts, between African and mixed ancestry cohorts and between white British and Asian ancestry cohorts.

We did not observe any significant heterogeneity across ancestries (genetic correlation estimate was not significantly different from 1) for waist-hip ratio (**Figure 5e and Supplementary Table 17**). Non-estimable cross-ancestry genetic correlation was observed when using African ancestry cohort (NA in **Figure 5e**) for which SNP-based heritability estimate was not significantly different from zero (**Figure 4 and Supplementary Table 17**).

## Discussion

We propose a novel method that provides unbiased estimates of ancestry-specific SNP-based heritability and cross-ancestry genetic correlations. This is possible because the proposed method correctly account for ancestry-specific genetic architectures or ancestry-specific heritability models. Our method provides a tool to dissect the ancestry-specific genetic architecture of a complex trait and can inform how genetic variance and covariance change across populations and ancestries. By using a meta-analysis across multiple ancestry groups^25,26^ based on unbiased estimates of ancestry-specific heritability and cross-ancestry genetic correlations, we hope the current ancestry disparity and study bias in GWAS^5,8^ can be reduced.

We investigated and found optimal *α* values for multiple ancestry groups, i.e. white British, other Europeans, Asian, African, and mixed ancestry cohorts, using six anthropometric traits from the UK Biobank. Interestingly, *α* values are distinct and dynamically distributed across ancestries even for the same complex trait, that is the relationship between the causal effects and allele frequency of the causal variants varies across ancestries. Per-allele effect size can be dynamically distributed, depending on genetic and environmental background or any unknown ancestry-specific factors. For example, if there are epistatic or interaction effects, selections on multiple loci can vary allele frequency, depending on per-allele effect sizes of loci. Moreover, SNP effects may reflect the level of association with causal variants such that per-allele effect size can be linearly correlated with allele frequency. Given our observation, it is clear that the heritability model should properly account for such diverse genetic architectures.

It was observed that the GCTA-*α* model outperformed LDAK-thni-*α* model for a more homogeneous population, such as white British, other Europeans or Asian ancestry cohort **(Supplementary Table 8).** For a less homogenous population such as mixed ancestry cohort, the LDAK-thin-*α* model was better than the GCTA-*α* model, implying that the choice of GCTA-*α* or LDAK-thin-*α* model might depend on the homogeneity of the population. It is noted that we used HapMap3 SNPs that have already excluded many redundant variants. A further study may be required to assess the performance of GCTA-*α* and LDAK-thin-*α* models with 1KG SNPs and other ancestry grouping, which is beyond the scope of this study.

The estimated SNP-based heritability of BMI in the white British ancestry cohort was not much different from previous studies^3,27^, although our estimate was slightly lower probably because of using different *α* values. For the same reason, our estimate for African ancestry cohort was slightly higher than previous study^3^. For standing height, we observed that the estimated SNP-based heritability in African ancestry cohort was significantly lower than other ancestry groups, which agreed with previous studies^3,28^. This is probably due to the fact that African-specific causal variants are less tagged by the common SNPs or environmental effects are relatively large, compared to other ancestries, which requires further investigations. For other traits, our estimates were approximately agreed with a previous study^27,29,30^ although using different *α* values.

Cross-ancestry genetic correlations can provide crucial information in cross-ancestry GWAS and cross-ancestry polygenic risk score prediction. We showed a significant genetic heterogeneity for obesity traits (BMI, weight and waist and hip circumferences) between African and European ancestry samples. For height, there was a significant genetic heterogeneity between African and Asian ancestry samples. However, without considering ancestry-specific *α* values (i.e. GRM 4 from equation 5), the findings were changed and could be over-interpreted, e.g. there was an additionally significant genetic heterogeneity between African and European ancestry cohorts (**Supplementary Figure 5**), which agreed with Guo et al.^3^ who used the same method as in GRM4 (equation 5).

LDSC based on GWAS summary statistics is computationally efficient and has been widely used in a single ancestry group (dominantly used in Europeans) in the estimation of SNP-based heritability and genetic correlations between complex traits^31^. Similar to LDSC, GWAS summary statistics-based cross-ancestry analyses (Popcorn)^2^ has been used by several studies^11–13^, to estimate cross-ancestry genetic correlation between European and east Asian. However, as shown in the results from Brown et al.^2^ and our simulation, the method can be biased when the genetic architecture of a complex traits is diverse (i.e. when *α* values vary) across ancestries. It is also noted that cross-ancestry meta GWAS may require unbiased estimates of cross-ancestry genetic correlations^25,26,32^.

There are several limitations in this study. Firstly, we used the GCTA-*α* model only in the real data analysis, assuming that all SNPs contributed equally to the heritability estimation^21,33^. We did not use the LDAK-thin-*α* model that required to prune SNPs within each ancestry group, which could substantially reduce the number of common SNPs between two ancestry groups in cross-ancestry genetic correlation analyses. Secondly, we did not consider MAF stratified, LDMS, or baseline model^34–36^ in estimating cross-ancestry genetic correlations because it was developed to estimate SNP-based heritability, but not suitably designed to estimate genetic correlations. Furthermore, it is required to fit multiple random effects (i.e. multiple GRMs), which is not computationally efficient. Nevertheless, these advanced models can improve the estimations of ancestry-specific SNP-based heritability and cross-ancestry genetic correlation. Thirdly, we estimated optimal scale factors (*α*) with a moderate sample size especially for Asian or African ancestry cohorts, resulting in a relatively flat curve of ΔAIC values. A further study is required to estimate more reliable *α* for Asian or African ancestry cohorts with a larger sample size.

In conclusion, we present a method to construct a GRM that can correctly account for the relationship between ancestry-specific allele frequencies and ancestry-specific causal effects. As the result, our method can provide unbiased estimates of ancestry-specific SNP-based heritability and cross-ancestry genetic correlation. By applying our proposed method to anthropometric traits, we found that obesity is a genetically heterogenous trait for African and European ancestry cohorts, while human height is a genetically heterogenous trait between African and Asian ancestry cohorts.

## Methods

### Ethics statement

We used data from the UK Biobank (https://www.ukbiobank.ac.uk), the scientific protocol of which has been reviewed and approved by the North West Multi-center Research Ethics Committee, National Information Governance Board for Health & Social Care, and Community Health Index Advisory Group. UK Biobank has obtained informed consent from all participants. Our access to the UK Biobank data was under the reference number 14575. The research ethics approval of this study has been obtained from the University of South Australia Human Research Ethics Committee.

### Statistical model

Our main aim is to unbiasedly estimate cross-ancestry genetic correlation for a complex trait, using common SNPs presented for both populations. We use a bivariate linear mixed model (LMM) to estimate SNP-based heritability and cross-ancestry genetic correlation, using GWAS data comprising multiple ancestry groups. In the model, a vector of phenotypic observations for each ancestry group can be decomposed into fixed effects, random genetic effects and residuals. The LMM can be written as,

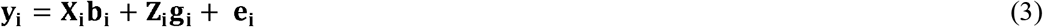

where **y_i_** is the vector of phenotypic observations, **b_i_** is the vector of fixed (environmental) effect with the incidence matrix **X_i_**, **g_i_** is the vector of random additive genetic effects with the incidence matrix **Z_i_** and **e_i_** is the vector of residual effects for the *i^th^* ancestry group (i = 1 and 2).

The random effects, **g_i_** and **e_i_**, are assumed to be normally distributed, i.e. 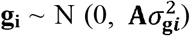 and 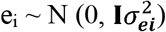. The variance covariance matrix of observed phenotypes can be written as

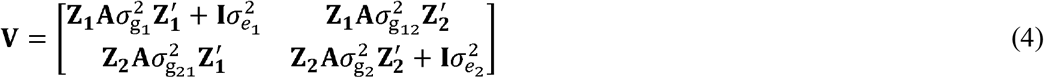

where **A** is the genomic relationship matrix (GRM)^19,20,37^, which can be estimated based on the genome-wide SNP information, and **I** is an identity matrix. The terms, 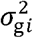 and 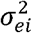, indicate the genetic and residual variance of the trait for the *i*^th^ ancestry group, and 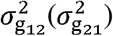 is the genetic covariances between the two ancestry groups. Note that it is not required to model residual correlation in **V** because there are no multiple phenotypic measures for any individual (no common residual effects shared between two ancestry groups).

### The variance of random additive genetic effects

Assuming that causal variants are in linkage equilibrium and that the phenotypic variance is *var*(*y*)=1, the heritability can be written as

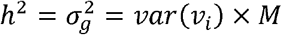

where *var*(*v_i_*) is the genetic variance of the *i^th^* causal variant and M is the number of causal variant. The genetic variance at the *i^th^* genetic variant can be written as

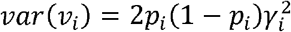

where *γ_i_* = *β_i_* are causal effects of the *i^th^* variant if we do not consider the relationship between *β_i_* and *p_i_*, following Falconer and Mackay^18^.

When considering the relationship between *β_i_* and *p_i_*, *γ_i_* can be reparameterised as *γ_i_* = *β_i_* × [2*p_i_*(1 – *p_i_*)]^*α*^ as suggested by previous studies^15,16,22^. This shows that, although *β_i_* is consistent, *γ_i_* can differ across ancestry groups that have different *p_i_* and/or *α*. Therefore, as shown in eq. (1), the genetic variance at the *i^th^* genetic variant can be rewritten as

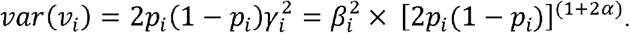

Assuming that the expectation of *β_i_* is E(*β_i_*) = 0, the expectation of *var*(*v_i_*) can be expressed as

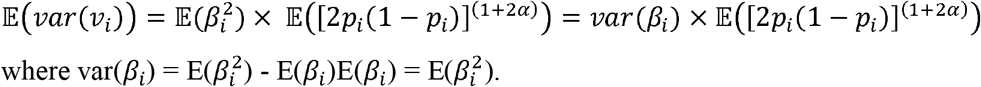

This shows 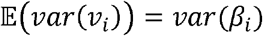 when using *α* = −0.5 (i.e. the widely used assumption of constant variance across different MAF spectrum).

However, with various factors (selection, interaction, linkage disequilibrium, population stratification and so on), optimal *α* values vary across populations^16,22^. Therefore, although per-allele effect size (*β_i_*) is constant, the actual effect, *γ_i_* = *β_i_* × (2*p_i_*(1 – *p_i_*)^*α*^), can be dynamically distributed, depending on ancestry-specific factors. What we aim to estimate is *cor*(**β**_*k*_, **β**_*l*_), the correlation between per-allele effect sizes for SNPs of the *k^th^* and *l^th^* ancestry groups, which is different from *cor*(**γ**_*k*_, **γ**_*l*_).

### Genomic Relationship Matrix (GRM)

GRM is a kernel matrix and can be normalized with a popular form of

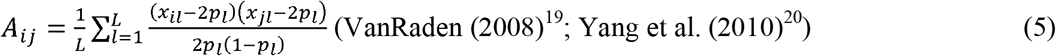

or

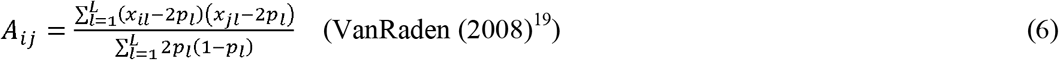

where *A_ij_* is the genomic relationship between the *i*^th^ and *j*^th^ individuals, *L* is the total number of SNPs, *p_l_* is the reference allele frequency at the *l*th SNP, and *x_il_* is the genotype coefficient of the *i*^th^ individual at the *l*^th^ SNP.

Speed et al.^15,16^ generalised these forms, introducing a scale factor that can determine the genetic architecture of a complex trait (aka heritability model). The generalised form is

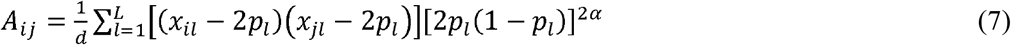

where *α* is a scale factor, *d* is the expected diagonals, 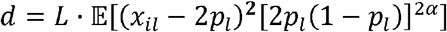. When *α* = −0.5, equation. (5) is equivalent to equation (3), and when *α* = 0, equation (5) becomes equivalent to equation (4). Note that each SNP can be weighted according to the LD structure (LDAK or LDAK-thin) if this weighting scheme better fits with the genetic architecture, which can be assessed by a model comparison^16^.

However, these equations (5, 6 and 7) do not account for correlation between *x_il_* and the estimated mean, i.e., 2*p_l_*, which can cause biased estimates of genomic relationships. With 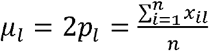 and *α* = −0.5, the diagonals can be expressed as

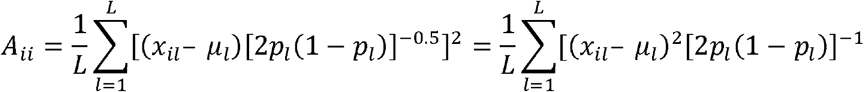

where the expectation of the first term can be rewritten as

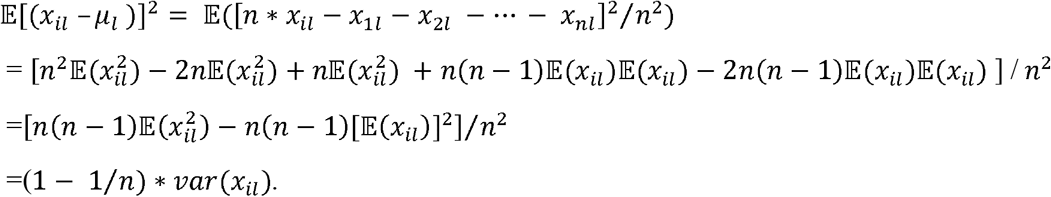

With 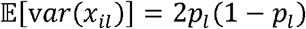, the diagonals from equation (5) or (7) can be expressed as

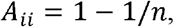

which is deviated from 1 by a factor 1/n. Without loss of generality, the biased factor with any *α* values can be written as

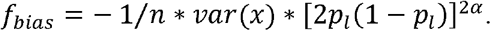

In a similar manner, the off diagonals can be written as

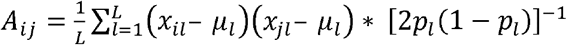

where the expectation of the first term can be rewritten as

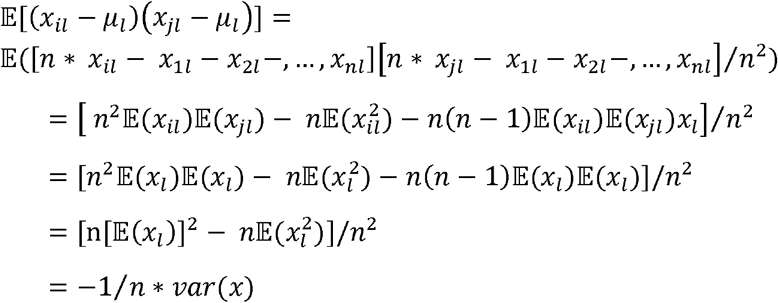

With 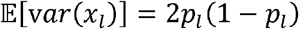, the off diagonals from equation (5) or (7) can be expressed as

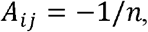

which is deviated from 0 by a factor 1/n. The biased factor with any *α* values is the same as in the diagonals, i.e. *f_bias_* = – 1/*n* * *var*(*x*) * [2*p_l_*(1 – *p_l_*)]^2*α*^.

Therefore, a revised formula, considering *f*_bias_, should be

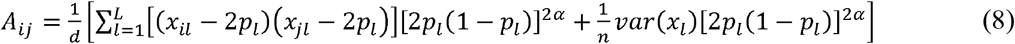

where *d* is redefined as 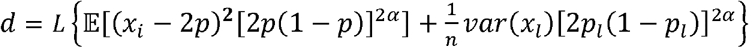. It is noted that with a sufficient sample size (> 1,000), the difference between equation (7) and (8) is negligible.

Furthermore, 2*p_l_*(1 – *p_l_*) is the expectation of var(*x*). Unless the samples are completely homogenous, the expectation is an approximation of var(*x*). So, var(*x*) should be used instead of the expectation 2*p*(1-*p*). Therefore, the formula can be rewritten as

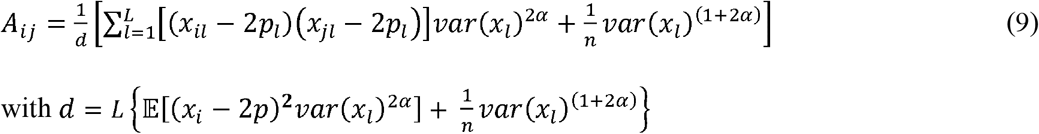

### GRM for cross-ancestry genetic analyses

Yang et al.^4^ proposed a GRM method that uses ancestry-specific allele frequency to be applied to cross-ancestry genetic analyses, which can be expressed as

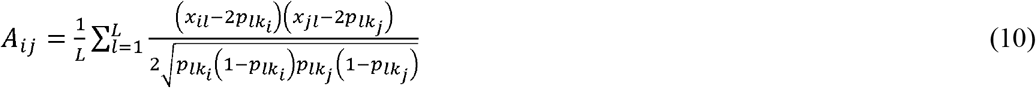

where *p_lk_i__* and *p_lk_j__* are the allele frequencies at the *l^th^* SNP estimated in the *k_i_* and *k_j_* th ancestry groups to which the *i^th^* and *j^th^* individuals belongs. When estimating cross-ancestry genetic correlation for Attention Deficit Hyperactivity Disorder (ADHD) between European and East Asian (Han Chinese), Yang et al.^4^ considered the standard scale factor for both ancestry groups (*α* = −0.5 in white European and East Asian). Similarly, Guo et al.^3^ also used equation (10) to estimate cross-ancestry genetic correlation between white British and African ancestry cohorts, assuming *α* value was constant across these ancestry groups (*α* = −0.5).

It is intuitive that *α* value can be dynamically changed across ancestry groups because of genetic drift, natural selection, and various selection pressures. Combined with a revised formula derived above (equation (9)), a novel GRM equation for cross-ancestry genetic analyses, which accounts for ancestry-specific *α* and ancestry-specific allele frequency, can be proposed as

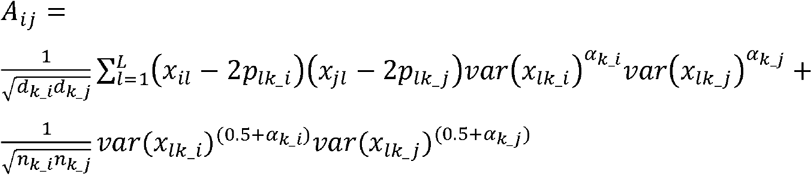

where *x_il_* and *x_jl_* are SNP genotypes for the *i^th^* and *j^th^* individuals at the *l*^th^ SNP, *p_lk_i_* and *p_lk_j_* are the allele frequencies at the *l*th SNP estimated in the two ancestry groups, *k_i* and *k_j*, to which the *i^th^* and *j^th^* individuals belongs, and *α_k_i_* and *α_k_j_* are the scale factors for the two ancestry groups, *x_lk_i_* and *x_lk_j_* are all individual genotypes at the lth SNP in the two ancestry groups, *n_k_i_* and *n_k_j_* are the number of individuals of the two ancestry groups, and *d_k_i_* is the expectation of the diagonals, 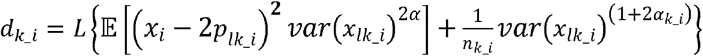.

### Data source and quality control

We tested this proposed method in real genotypic and phenotypic data obtained from the second release of the UK Biobank (https://www.ukbiobank.ac.uk/). The genotypic data were imputed based on Haplotype Reference Consortium reference panel^38^. The UK Biobank data comprised 488,377 participants and 92,693,895 SNPs. The participants were grouped into five ancestry groups (white British, other European, Asian, African, and mixed ancestry cohorts) according to their genetic ancestry estimated from a principal component analysis based on the genome-wide SNP infromation^17^. Mixed ancestry cohort includes individuals from Asian ancestry, some white and black African, some white and black Caribbean and those individuals assigned as other ancestry groups in UK biobank (**Supplementary Figure1**). We did not include individuals who do not know their ancestry and who prefer not to answer (UK Biobank codes are −1 and −6). Gender mismatch and sex chromosome aneuploidy were also excluded during the quality control (QC) process.

We performed additional stringent QC in each of the ancestry groups. The QC criteria include an INFO score (an imputation reliability) ≥0.6^39–41^, SNP missingness <0.05, minor allele frequency (MAF) >0.01, Hardy–Weinberg equilibrium p-value > 10^-04^. We also excluded individuals outside ±6 SD of the population mean for first and second ancestry principal components. Individuals with genetic relatedness ≥0.05 were excluded from each ancestry group using PLINK^42^. In the analysis, we retained HapMap3 SNPs only as these are high in quality and well calibrated to dissect genetic architecture of complex traits^43,44^.

The initial sample size was 430301, 29023, 7449, 7647 and 16615 for white British, other European, Asian, African, and mixed ancestry cohorts, respectively. After quality control, the cleaned data includes 288837, 26457, 6199, 6179 and 11797 participants, and the total number of SNP was 1154490, 1148504, 939512, 729534 and 513362 for white British, other European, Asian, African, and mixed ancestry cohorts. For white British ancestry cohort, we randomly selected 30000 from the QCed 288837 individuals for estimating cross-ancestry genetic correlations because it is computationally feasible in multiple analyses paring with other ancestry groups.

### Phenotype Simulation

The phenotypes of each ancestry group were simulated using a bivariate linear mixed model (equation 3, i.e. **y_i_** = **X_i_b_i_** + **Z_i_g_i_** + **e_i_**). In the simulation, 1000 individuals and 500000 SNPs were randomly chosen for each ancestry group. To simulate phenotypes, we randomly selected 1000 SNPs that were common between the two ancestry groups and assigned causal effects to them. According to equation (4), a multivariate normal distribution was used to draw causal effects of the 1000 SNPs with mean 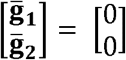, and genetic covariance matrix as 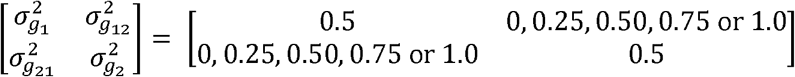. According to equation (1), the causal effects of the *i^th^* SNPs were scaled by actual variance [*var*(*x_i_*)]^*α*^, where *p_i_* is the reference allele frequency, noting that *α* and *p_i_* can vary between the two ancestry groups. Individual genetic values (i.e. polygenic risk scores) are the sum of their genotype coefficients of the 1000 causal SNPs, weighted by the causal effects. The simulated phenotypes were generated as the summation of the true genetic values and the residual effects (equation 4) which were obtained from a multivariate normal distribution with mean 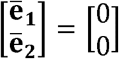 and the residual covariance matrix 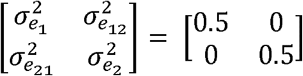. Hence, the true heritability was set as 0.5 for both ancestry groups in the simulation based on the bivariate linear mixed model.

To validate the proposed method of GRM estimation, we considered scenarios with the number of causal SNPs 100, 1000, 10000 and 100000. Phenotypic data were simulated based on standard *α* and estimated *α* during scaling of random causal effect (equation 1). The simulation process was performed using R, PLINK^42^ and MTG2^45^. The biasedness of estimates was assessed by Wald test.

### Determining scale factor (*α*) across ancestries for LDAK-thin-α and GCTA-α model

We analysed six different anthropometric traits (BMI, standing height, waist circumference, hip circumference, waist hip ratio and weight) from the UK Biobank across different ancestries (white British, other European, Asian, African, and mixed ancestry cohorts). These traits were adjusted for demographic variables, UK biobank assessment centre, genotype measurement batch and population structure measured by the first 10 principal components (PCs)^27,46^. Demographic variable includes sex, year of birth, education, and Townsend deprivation index. Information of educational qualifications converted to education levels (years) for all the UK Biobank individuals^47^.

GCTA-*α* and LDAK-thin-*α* models^22^ were used to determine optimal *α* values for each of the ancestry groups. We considered various *α* values, e.g. *α* = −1, −0.875, −0.75, −0.675, −0.5, −0.375, −0.25, −0.125, 0 and 0.125, following the approach of Speed et al^16^. All GRMs with various *α* values were estimated using LDAK software^15^, which set an equal weight to all SNPs in GCTA-*α* model and different weights to SNPs according to their LD scores in LDAK-thin-*α* model. Using GRMs, SNP-based heritabilities were estimated for six anthropometric traits, using a multivariate linear mixed model^45^ that fit six anthropometric traits simultaneously. Note that in the multivariate model, we treated the six traits independent without considering residual and genetic correlations between traits. This was because the optimal alpha value should be trait-specific as our main analysis was to estimate trait-specific cross-ancestry genetic correlation, i.e. equal weights from all six traits to obtain the optimal *α* value for each ancestry group. Finally, we identified optimal *α* that gives lowest AIC values, i.e. AIC = 2*k* – 2*ln*(*L*) where ln(*L*) is the logarithm of the maximum likelihood from the model and k is the number of parameters.

### Genetic correlation estimation using existing methods

Bivariate GREML^11,23,48,49^ analyses were used to estimate heritability and cross-ancestry genetic correlation. In the analyses, we used four existing GRM methods (**Table 1)** to assess their performance, compared to the proposed method using simulated phenotypes, using PLINK^42^ (*-make-grm-gz* for command GRM1 and GRM2) and GCTA^21^ software (*-sub-popu* command for GRM3 and GRM4). The distribution of diagonal elements and off-diagonal elements across ancestries represented in **Supplementary Figure 6-10** (using α =-0.5 and estimated in PLINK2). We also used Popcorn^2,11^, that could be based on GWAS summary statistics. For these methods, we calculated empirical SE and its 95% confidence interval (CI) over 500 replicates to assess the unbiased estimation of the methods for each combination of ancestry pairs.

## Acknowledgements

This research is supported by the Australian Research Council (DP190100766) and the software development is supported by Cooperative Research Program for Agriculture Science and Technology Development (PJ0160992021) from the Rural Development Administration, Republic of Korea. We are grateful to Professors Peter Visscher and Doug Speed for their constructive criticisms and comments on the manuscript. We thank the staff and participants of the UK Biobank for their important contributions. The analyses were performed using computational resources provided by the Australian Government through Gadi under the National Computational Merit Allocation Scheme (NCMAS), and HPCs (Tango and Statgen server) managed by UniSA IT.

## Web resources

The genotype and phenotype data of the UK Biobank can be accessed through procedures described on its webpage (https://www.ukbiobank.ac.uk/). Simulated data used in this paper can be obtained from the authors upon request.

MTG2, https://sites.google.com/site/honglee0707/mtg2

GCTA, https://cnsgenomics.com/software/gcta/#Download

LDAK, http://dougspeed.com/ldak/

PLINK, https://www.cog-genomics.org/plink/

Popcorn, https://github.com/brielin/popcorn

## Code availability

The source code for MTG2 and example code along with related files for estimating GRM in combined ancestries using MTG2 can be accessed from https://sites.google.com/site/honglee0707/mtg2

