## Supplementary information for "A novel method for an unbiased estimate of cross-ancestry genetic correlation using individual-level data"

**Supplementary Table 1: Theoretical bias factors and estimated genomic relationships when using the standard scaling (*α* = -0.5).**

| **Number of random population samples (n)** | **Bias factor** | **Mean (**±**SD) of off-diagonals in estimated GRM** | |  | **Mean (**±**SD) of diagonals in estimated GRM** | |
| --- | --- | --- | --- | --- | --- | --- |
|  |  | **Existing method (equation 5)** | **Proposed method (equation 2)** |  | **Existing method (equation 5)** | **Proposed method (equation 2)** |
| 10 | -0.1 | -0.0961±0.0035 | -0.0094±0.0044 |  | 0.8592±0.0032 | 1.0±0.0037 |
| 100 | -0.01 | -0.0101±0.0045 | 0.0000±0.0044 |  | 0.9936±0.0127 | 1.0±0.0112 |
| 1000 | -0.001 | -0.0010±0.0046 | 0.0000±0.0046 |  | 0.9997±0.0143 | 1.0±0.0138 |
| 10000 | -0.0001 | -0.0001±0.0046 | 0.0000±0.0046 |  | 1.0003±0.0135 | 1.0±0.0132 |

GRM was estimated for white British based on 500,000 SNPs with various sample sizes. Diagonals and off-diagonals are biasedly estimated when using existing method (i.e. equation 5), noting that the bias factor is $f_{bias}={-1}/{n*var(x)}*\left[ 2p_{l}\left( 1-p_{l} \right) \right]^{2\alpha}=-1/n$ (with *α* = -0.50) (see equations 5-8 in Methods). When correcting the bias factor, the proposed method (i.e., equation 2 or 8) provides unbiased estimates.

**Supplementary Table 2: Slightly biased estimates of SNP-based heritability when using the expected variance in scaling genotypic coefficients in constructing GRM.**

|  | **Using actual variance (var(x)) in scaling genotypic coefficients to estimate GRM** | |  | **Using expected variance (2p(1-p)) in scaling genotypic coefficients to estimate GRM** | |
| --- | --- | --- | --- | --- | --- |
| **Ancestry** | **Estimated** $\boldsymbol{h}^{\boldsymbol{2}}$**(**±SE) | **P-value** |  | **Estimated** $\boldsymbol{h}^{\boldsymbol{2}}$**(**±SE) | **P-value** |
| British | 0.479±0.013 | 0.1062 |  | 0.484±0.013 | 0.2184 |
| Asian | 0.483±0.013 | 0.1910 |  | 0.474±0.013 | 0.0455 |
| Africa | 0.508±0.012 | 0.5050 |  | 0.465±0.012 | 0.0035 |
| Mixed ancestry | 0.489±0.009 | 0.2216 |  | 0.476±0.010 | 0.0163 |

The true heritability was 0.50. Mean estimate and SE were calculated over 500 replicates. P-values were from Wald test (null hypothesis of estimated $h^{2}$= 0.50). In simulation, the causal SNP effects were scaled by actual variance (see method section for details). We observed biased heritability estimates (red coloured) in Asian, African and mixed ancestries when the estimation was based on the expected variance instead of the actual variance.

**Supplementary Table 3: Inferring scaling factors in white British ancestry cohort for LDAK-thin-α and GCTA-*α* models.**

| **Log-likelihood of White British ancestry cohort (n=30000), LDAK-thin-*α* model** | | | | | | | |
| --- | --- | --- | --- | --- | --- | --- | --- |
| **Traits** | ***α*= -0.625** | ***α*= -0.50** | ***α*= -0.375** | ***α*= -0.25** | ***α*= -0.125** | ***α* =0** | ***α* =0.125** |
| **BMI** | -14648.386 | -14641.042 | -14638.387 | -14638.168 | -14639.145 | -14640.688 | -14642.484 |
| **Standing Height** | -13981.958 | -13939.456 | -13919.496 | -13912.319 | -13912.013 | -13915.356 | -13920.646 |
| **Waist circumference** | -14701.715 | -14697.099 | -14695.797 | -14696.171 | -14697.357 | -14698.923 | -14700.655 |
| **Hip circumference** | -14681.115 | -14674.272 | -14671.691 | -14671.314 | -14672.012 | -14673.214 | -14674.641 |
| **Waist-hip ratio** | -14746.071 | -14741.714 | -14739.861 | -14739.388 | -14739.669 | -14740.370 | -14741.302 |
| **Weight** | -14587.158 | -14578.471 | -14575.913 | -14576.464 | -14578.471 | -14581.111 | -14583.984 |
| **Multivariate model^a^** | **-87346.403** | **-87272.054** | **-87241.145** | **-87233.824** | **-87238.667** | **-87249.662** | **-87263.712** |
| **AIC^b^** | **174704.806** | **174556.108** | **174494.29** | **174479.648** | **174489.334** | **174511.324** | **174539.424** |
| **ΔAIC^c^** | **225.158** | **76.46** | **14.642** | **0** | **9.686** | **31.676** | **59.776** |
| **Log-likelihood of White British ancestry cohort (n=30000), GCTA-*α* model** | | | | | | | |
| **Traits** | ***α*= -0.625** | ***α*= -0.50** | ***α*= -0.375** | ***α*= -0.25** | ***α*= -0.125** | ***α* =0** | ***α* =0.125** |
| **BMI** | -14646.809 | -14637.704 | -14633.573 | -14632.214 | -14632.345 | -14633.272 | -14634.626 |
| **Standing Height** | -13916.334 | -13869.658 | -13846.069 | -13836.442 | -13834.900 | -13838.006 | -13843.798 |
| **Waist circumference** | -14703.271 | -14697.264 | -14694.827 | -14694.348 | -14694.914 | -14696.035 | -14697.451 |
| **Hip circumference** | -14680.958 | -14673.030 | -14669.419 | -14668.202 | -14668.272 | -14669.032 | -14670.163 |
| **Waist-hip ratio** | -14744.532 | -14739.444 | -14737.009 | -14736.091 | -14736.032 | -14736.471 | -14737.202 |
| **Weight** | -14579.028 | -14568.363 | -14564.171 | -14563.515 | -14564.692 | -14566.794 | -14569.344 |
| **Multivariate model^a^** | **-87270.932** | **-87185.463** | **-87145.068** | **-87130.812** | **-87131.155** | **-87139.61** | **-87152.584** |
| **AIC^b^** | **174553.864** | **174382.926** | **174302.136** | **174273.624** | **174274.31** | **174291.22** | **174317.168** |
| **ΔAIC^c^** | **280.24** | **109.302** | **28.512** | **0** | **0.686** | **17.596** | **43.544** |

^a^Multivariate linear mixed model was used to get the log-likelihood of the scaling factor where residual and genetic correlations between traits were fixed as zero, i.e. the log-likelihood of this multivariate linear mixed model is the sum of log-likelihood values from the trait-specific analyses. ^b^Akaike Information Criterion $\left( \mathrm{AIC} \right)=2k-2ln(L)$ where $2\ln(L)$ is the logarithm of the maximum likelihood from the model and k is the number of model parameters in the model. ^c^ΔAIC = AIC – AIC of the best model with the optimal *α*. The best model is red highlighted.

**Supplementary Table 4: Inferring scaling factors in other European ancestry cohort for LDAK-thin-*α* and GCTA-*α* models.**

| **Log-likelihood of other European ancestry cohort (n=26457), LDAK-thin-*α* model** | | | | | | | |
| --- | --- | --- | --- | --- | --- | --- | --- |
| **Traits** | ***α*= -0.625** | ***α*= -0.50** | ***α*= -0.375** | ***α*= -0.25** | ***α*= -0.125** | ***α* =0** | ***α* =0.125** |
| **BMI** | -12825.531 | -12817.869 | -12814.507 | -12813.513 | -12813.782 | -12814.721 | -12816.014 |
| **Standing Height** | -12314.739 | -12288.168 | -12277.591 | -12275.302 | -12277.133 | -12281.057 | -12286.098 |
| **Waist circumference** | -12894.232 | -12889.131 | -12886.667 | -12885.639 | -12885.384 | -12885.563 | -12885.998 |
| **Hip circumference** | -12875.624 | -12868.142 | -12864.204 | -12862.319 | -12861.599 | -12861.555 | -12861.916 |
| **Waist-hip ratio** | -12918.195 | -12914.027 | -12911.747 | -12910.574 | -12910.057 | -12909.947 | -12910.099 |
| **Weight** | -12816.469 | -12806.169 | -12800.980 | -12798.712 | -12798.070 | -12798.346 | -12799.151 |
| **Multivariate model^a^** | **-76644.790** | **-76583.506** | **-76555.696** | **-76546.059** | **-76546.025** | **-76551.189** | **-76559.276** |
| **AIC^b^** | **153301.58** | **153179.012** | **153123.392** | **153104.118** | **153104.05** | **153114.378** | **153130.552** |
| **ΔAIC^c^** | **197.53** | **74.962** | **19.342** | **0.068** | **0** | **10.328** | **26.502** |
| **Log-likelihood of other European ancestry cohort (n=26457), GCTA-*α* model** | | | | | | | |
| **Traits** | ***α*= -0.625** | ***α*= -0.50** | ***α*= -0.375** | ***α*= -0.25** | ***α*= -0.125** | ***α* =0** | ***α* =0.125** |
| **BMI** | -12822.089 | -12814.182 | -12810.608 | -12809.513 | -12809.781 | -12810.791 | -12812.197 |
| **Standing Height** | -12282.774 | -12255.861 | -12243.939 | -12240.204 | -12240.845 | -12243.901 | -12248.364 |
| **Waist circumference** | -12889.703 | -12884.144 | -12881.440 | -12880.387 | -12880.267 | -12880.679 | -12881.398 |
| **Hip circumference** | -12870.057 | -12862.691 | -12858.841 | -12857.059 | -12856.466 | -12856.567 | -12857.083 |
| **Waist-hip ratio** | -12916.542 | -12911.875 | -12909.306 | -12908.041 | -12907.572 | -12907.594 | -12907.922 |
| **Weight** | -12809.056 | -12798.907 | -12793.705 | -12791.405 | -12790.771 | -12791.108 | -12792.022 |
| **Multivariate model^a^** | **-76590.221** | **-76527.660** | **-76497.839** | **-76486.609** | **-76485.702** | **-76490.640** | **-76498.986** |
| **AIC^b^** | **153192.442** | **153067.32** | **153007.678** | **152985.218** | **152983.404** | **152993.28** | **153009.972** |
| **ΔAIC^c^** | **209.038** | **83.916** | **24.274** | **1.814** | **0** | **9.876** | **26.568** |

^a^Multivariate linear mixed model was used to get the log-likelihood of the scaling factor where residual and genetic correlations between traits were fixed as zero, i.e. the log-likelihood of this multivariate linear mixed model is the sum of log-likelihood values from the trait-specific analyses. ^b^Akaike Information Criterion $\left( \mathrm{AIC} \right)=2k-2ln(L)$ where 2*ln*(*L*) is the logarithm of the maximum likelihood from the model and k is the number of model parameters in the model. ^c^ΔAIC = AIC – AIC of the best model with the optimal *α*. The best model is red highlighted.

**Supplementary Table 5: Inferring scaling factors in Asian ancestry cohort for LDAK-thin-*α* and GCTA-*α* models**

| **Log-likelihood of Asian ancestry cohort (n=6199), LDAK-thin-*α* model** | | | | | | | | | | |
| --- | --- | --- | --- | --- | --- | --- | --- | --- | --- | --- |
| **Traits** | ***α*= -1** | ***α*= -0.875** | ***α*= -0.75** | ***α*= -0.625** | ***α*= -0.50** | ***α*= -0.375** | ***α*= -0.25** | ***α*= -0.125** | ***α*= 0** | ***α*= 0.125** |
| **BMI** | -2850.005 | -2848.154 | -2847.217 | -2847.138 | -2847.531 | -2848.068 | -2848.596 | -2849.070 | -2849.483 | -2849.846 |
| **Standing Height** | -2841.233 | -2832.923 | -2826.611 | -2823.219 | -2822.141 | -2822.336 | -2823.049 | -2823.885 | -2824.672 | -2825.355 |
| **Waist circumference** | -2902.485 | -2901.083 | -2900.065 | -2899.583 | -2899.510 | -2899.643 | -2899.849 | -2900.068 | -2900.271 | -2900.453 |
| **Hip circumference** | -2900.520 | -2900.467 | -2900.895 | -2901.583 | -2902.269 | -2902.831 | -2903.255 | -2903.571 | -2903.809 | -2903.994 |
| **Waist-hip ratio** | -2907.148 | -2906.052 | -2904.386 | -2902.869 | -2901.899 | -2901.465 | -2901.402 | -2901.555 | -2901.812 | -2902.113 |
| **Weight** | -2892.291 | -2889.713 | -2887.985 | -2887.286 | -2887.301 | -2887.651 | -2888.106 | -2888.568 | -2888.999 | -2889.392 |
| **Multivariate model^a^** | **-17293.682** | **-17278.392** | **-17267.159** | **-17261.678** | **-17260.651** | **-17261.994** | **-17264.257** | **-17266.717** | **-17269.046** | **-17271.253** |
| **AIC^b^** | **34599.364** | **34568.784** | **34546.318** | **34535.356** | **34533.302** | **34535.988** | **34540.514** | **34545.434** | **34550.092** | **34554.506** |
| **ΔAIC^c^** | **66.062** | **35.482** | **13.016** | **2.054** | **0** | **2.686** | **7.212** | **12.132** | **16.79** | **21.204** |
| **Log-likelihood of Asian ancestry cohort (n=6199), GCTA-*α* model** | | | | | | | | | | |
| **Traits** | ***α*= -1** | ***α*= -0.875** | ***α*= -0.75** | ***α*= -0.625** | ***α*= -0.50** | ***α*= -0.375** | ***α*= -0.25** | ***α*= -0.125** | ***α*= 0** | ***α*= 0.125** |
| **BMI** | -2848.455 | -2847.351 | -2847.214 | -2847.781 | -2848.609 | -2849.413 | -2850.099 | -2850.661 | -2851.126 | -2851.517 |
| **Standing Height** | -2836.662 | -2828.602 | -2823.144 | -2820.685 | -2820.319 | -2820.979 | -2821.974 | -2822.979 | -2823.877 | -2824.636 |
| **Waist circumference** | -2901.579 | -2900.588 | -2900.065 | -2899.999 | -2900.194 | -2900.474 | -2900.752 | -2900.999 | -2901.212 | -2901.394 |
| **Hip circumference** | -2898.808 | -2899.118 | -2900.006 | -2901.114 | -2902.113 | -2902.885 | -2903.446 | -2903.850 | -2904.145 | -2904.366 |
| **Waist-hip ratio** | -2907.145 | -2906.279 | -2905.106 | -2904.144 | -2903.606 | -2903.439 | -2903.509 | -2903.706 | -2903.956 | -2904.219 |
| **Weight** | -2889.802 | -2887.788 | -2886.776 | -2886.719 | -2887.201 | -2887.849 | -2888.489 | -2889.059 | -2889.557 | -2889.993 |
| **Multivariate model^a^** | **-17282.451** | **-17269.726** | **-17262.311** | **-17260.442** | **-17262.042** | **-17265.039** | **-17268.269** | **-17271.254** | **-17273.873** | **-17276.125** |
| **AIC^b^** | **34576.902** | **34551.452** | **34536.622** | **34532.884** | **34536.084** | **34542.078** | **34548.538** | **34554.508** | **34559.746** | **34564.25** |
| **ΔAIC^c^** | **44.018** | **18.568** | **3.738** | **0** | **3.2** | **9.194** | **15.654** | **21.624** | **26.862** | **31.366** |

^a^Multivariate linear mixed model was used to get the log-likelihood of the scaling factor where residual and genetic correlations between traits were fixed as zero, i.e. the log-likelihood of this multivariate linear mixed model is the sum of log-likelihood values from the trait-specific analyses. ^b^Akaike Information Criterion $\left( \mathrm{AIC} \right)=2k-2ln(L)$ where 2*ln*(*L*) is the logarithm of the maximum likelihood from the model and k is the number of model parameters in the model. ^c^ΔAIC = AIC – AIC of the best model with the optimal *α*. The best model is red highlighted.

**Supplementary Table 6: Inferring scaling factors in African ancestry cohort for LDAK-thin-*α* and GCTA-*α* models**

| **Log-likelihood of African ancestry cohort (n=6179), LDAK-thin-*α* model** | | | | | | | | | | |
| --- | --- | --- | --- | --- | --- | --- | --- | --- | --- | --- |
| **Traits** | ***α*= -1** | ***α*= -0.875** | ***α*= -0.75** | ***α*= -0.625** | ***α*= -0.50** | ***α*= -0.375** | ***α*= -0.25** | ***α*= -0.125** | ***α*= 0** | ***α*= 0.125** |
| **BMI** | -2927.518 | -2927.668 | -2928.253 | -2929.404 | -2930.789 | -2931.979 | -2932.837 | -2933.418 | -2933.816 | -2934.106 |
| **Standing Height** | -2937.988 | -2934.909 | -2931.289 | -2928.199 | -2926.395 | -2925.744 | -2925.730 | -2925.971 | -2926.284 | -2926.597 |
| **Waist circumference** | -2939.807 | -2939.515 | -2939.597 | -2940.323 | -2941.455 | -2942.544 | -2943.381 | -2943.977 | -2944.402 | -2944.721 |
| **Hip circumference** | -2936.153 | -2935.935 | -2936.247 | -2937.274 | -2938.686 | -2939.993 | -2940.985 | -2941.686 | -2942.186 | -2942.561 |
| **Waist-hip ratio** | -2949.119 | -2949.296 | -2949.528 | -2949.667 | -2949.053 | -2948.424 | -2947.938 | -2947.602 | -2947.379 | -2947.233 |
| **Weight** | -2935.594 | -2934.865 | -2934.475 | -2934.471 | -2935.076 | -2935.818 | -2936.457 | -2936.946 | -2937.317 | -2937.613 |
| **Multivariate model^a^** | **-17626.179** | **-17622.2** | **-17619.4** | **-17619.3** | **-17621.5** | **-17624.5** | **-17627.3** | **-17629.6** | **-17631.4** | **-17632.8** |
| **AIC^b^** | **35264.358** | **35256.38** | **35250.78** | **35250.68** | **35254.91** | **35261** | **35266.66** | **35271.2** | **35274.77** | **35277.66** |
| **ΔAIC^c^** | **13.682** | **5.7** | **0.102** | **0** | **4.232** | **10.328** | **15.98** | **20.524** | **24.092** | **26.986** |
| **Log-likelihood of African ancestry cohort (n=6179), GCTA-*α* model** | | | | | | | | | | |
| **Traits** | ***α*= -1** | ***α*= -0.875** | ***α*= -0.75** | ***α*= -0.625** | ***α*= -0.50** | ***α*= -0.375** | ***α*= -0.25** | ***α*= -0.125** | ***α*= 0** | ***α*= 0.125** |
| **BMI** | -2927.832 | -2928.195 | -2929.163 | -2930.665 | -2932.198 | -2933.397 | -2934.226 | -2934.791 | -2935.189 | -2935.490 |
| **Standing Height** | -2937.849 | -2934.538 | -2930.892 | -2928.063 | -2926.565 | -2926.078 | -2926.105 | -2926.334 | -2926.617 | -2926.899 |
| **Waist circumference** | -2939.803 | -2939.923 | -2940.618 | -2941.887 | -2943.268 | -2944.381 | -2945.165 | -2945.704 | -2946.089 | -2946.380 |
| **Hip circumference** | -2937.302 | -2937.341 | -2938.024 | -2939.333 | -2940.798 | -2942.017 | -2942.904 | -2943.531 | -2943.989 | -2944.343 |
| **Waist-hip ratio** | -2949.289 | -2949.363 | -2949.555 | -2949.763 | -2949.447 | -2949.134 | -2948.896 | -2948.734 | -2948.629 | -2948.563 |
| **Weight** | -2936.051 | -2935.517 | -2935.394 | -2935.875 | -2936.695 | -2937.496 | -2938.133 | -2938.616 | -2938.991 | -2939.297 |
| **Multivariate model^a^** | **-17628.126** | **-17624.9** | **-17623.6** | **-17625.6** | **-17629** | **-17632.5** | **-17635.4** | **-17637.7** | **-17639.5** | **-17640.97** |
| **AIC^b^** | **35268.252** | **35261.75** | **35259.29** | **35263.17** | **35269.94** | **35277.01** | **35282.86** | **35287.42** | **35291.01** | **35293.94** |
| **ΔAIC^c^** | **8.96** | **2.462** | **0** | **3.88** | **10.65** | **17.714** | **23.566** | **28.128** | **31.716** | **34.652** |

^a^Multivariate linear mixed model was used to get the log-likelihood of the scaling factor where residual and genetic correlations between traits were fixed as zero, i.e. the log-likelihood of this multivariate linear mixed model is the sum of log-likelihood values from the trait-specific analyses. ^b^Akaike Information Criterion $\left( \mathrm{AIC} \right)=2k-2ln(L)$ where 2ln(*L*) is the logarithm of the maximum likelihood from the model and k is the number of model parameters in the model. ^c^ΔAIC = AIC – AIC of the best model with the optimal *α*. The best model is red highlighted.

**Supplementary Table 7: Inferring scaling factors in mixed ancestry cohort for LDAK-thin-*α* and GCTA-*α* models**

| **Log-likelihood of Mixed ancestry cohort (n=11979), LDAK-thin-*α* model** | | | | | | | | | | |
| --- | --- | --- | --- | --- | --- | --- | --- | --- | --- | --- |
| **Traits** | ***α*= -1** | ***α*= -0.875** | ***α*= -0.75** | ***α*= -0.625** | ***α*= -0.50** | ***α*= -0.375** | ***α*= -0.25** | ***α*= -0.125** | ***α*= 0** | ***α*= 0.125** |
| **BMI** | -5588.259 | -5585.895 | -5588.577 | -5594.217 | -5599.984 | -5604.549 | -5607.815 | -5610.084 | -5611.664 | -5612.583 |
| **Standing Height** | -5596.967 | -5580.956 | -5569.985 | -5565.634 | -5565.784 | -5567.876 | -5570.469 | -5573.009 | -5575.333 | -5577.420 |
| **Waist circumference** | -5667.104 | -5664.900 | -5665.788 | -5668.692 | -5671.871 | -5674.420 | -5676.215 | -5677.407 | -5678.176 | -5678.659 |
| **Hip circumference** | -5660.905 | -5659.907 | -5661.853 | -5665.352 | -5668.692 | -5671.190 | -5672.886 | -5674.006 | -5674.749 | -5675.251 |
| **Waist-hip ratio** | -5686.903 | -5684.125 | -5682.170 | -5681.575 | -5681.901 | -5682.573 | -5683.274 | -5683.889 | -5684.399 | -5684.811 |
| **Weight** | -5641.025 | -5636.521 | -5636.154 | -5638.865 | -5642.432 | -5645.546 | -5647.911 | -5649.638 | -5650.909 | -5651.872 |
| **Multivariate model^a^** | **-33841.163** | **-33812.304** | **-33804.527** | **-33814.335** | **-33830.664** | **-33846.154** | **-33858.57** | **-33868.033** | **-33875.23** | **-33880.596** |
| **AIC^b^** | **67694.326** | **67636.608** | **67621.054** | **67640.67** | **67673.328** | **67704.308** | **67729.14** | **67748.066** | **67762.46** | **67773.192** |
| **ΔAIC^c^** | **73.272** | **15.554** | **0** | **19.616** | **52.274** | **83.254** | **108.086** | **127.012** | **141.406** | **152.138** |
| **Log-likelihood of Mixed ancestry cohort (n= 11979), GCTA-*α* model** | | | | | | | | | | |
| **Traits** | ***α*= -1** | ***α*= -0.875** | ***α*= -0.75** | ***α*= -0.625** | ***α*= -0.50** | ***α*= -0.375** | ***α*= -0.25** | ***α*= -0.125** | ***α*= 0** | ***α*= 0.125** |
| **BMI** | -5590.849 | -5590.816 | -5594.943 | -5600.645 | -5605.712 | -5609.472 | -5612.091 | -5613.901 | -5615.172 | -5616.089 |
| **Standing Height** | -5593.909 | -5579.833 | -5571.487 | -5568.877 | -5569.532 | -5571.411 | -5573.556 | -5575.638 | -5577.569 | -5579.339 |
| **Waist circumference** | -5667.672 | -5667.488 | -5669.858 | -5673.188 | -5676.094 | -5678.174 | -5679.547 | -5680.424 | -5680.975 | -5681.317 |
| **Hip circumference** | -5661.356 | -5661.804 | -5664.812 | -5668.569 | -5671.682 | -5673.851 | -5675.278 | -5676.214 | -5676.843 | -5677.280 |
| **Waist-hip ratio** | -5687.339 | -5685.519 | -5684.519 | -5684.413 | -5684.776 | -5685.260 | -5685.714 | -5686.097 | -5686.408 | -5686.658 |
| **Weight** | -5641.291 | -5638.975 | -5640.409 | -5643.792 | -5647.192 | -5649.854 | -5651.777 | -5653.159 | -5654.184 | -5654.978 |
| **Multivariate model^a^** | **-33842.416** | **-33824.435** | **-33826.028** | **-33839.484** | **-33854.988** | **-33868.022** | **-33877.963** | **-33885.433** | **-33891.151** | **-33895.661** |
| **AIC^b^** | **67696.832** | **67660.87** | **67664.056** | **67690.968** | **67721.976** | **67748.044** | **67767.926** | **67782.866** | **67794.302** | **67803.322** |
| **ΔAIC^c^** | **35.962** | **0** | **3.186** | **30.098** | **61.106** | **87.174** | **107.056** | **121.996** | **133.432** | **142.452** |

^a^Multivariate linear mixed model was used to get the log-likelihood of the scaling factor where residual and genetic correlations between traits were fixed as zero, i.e. the log-likelihood of this multivariate linear mixed model is the sum of log-likelihood values from the trait-specific analyses. ^b^Akaike Information Criterion $\left( \mathrm{AIC} \right)=2k-2\ln(L)$ where 2*ln*(*L*) is the logarithm of the maximum likelihood from the model and k is the number of model parameters in the model. ^c^ΔAIC = AIC – AIC of the best model with the optimal *α*. The best model is red highlighted.

**Supplementary Table 8: Difference (AIC of LDAK-thin-*α* – AIC of GCTA-*α*) between two models**

| **Ancestry** | **Differences of Log-likelihood between of two model** | | | | | | | | | | |
| --- | --- | --- | --- | --- | --- | --- | --- | --- | --- | --- | --- |
|  | ***α*= -1** | ***α*= -0.875** | ***α*= -0.75** | ***α*= -0.625** | ***α*= -0.50** | ***α*= -0.375** | ***α*= -0.25** | ***α*= -0.125** | ***α*= 0** | ***α*= 0.125** | **Best *α* and model** |
| **White British** | - | - | - | 150.942 | 173.182 | 192.154 | 206.024 | 215.024 | 220.104 | 222.256 | **-0.25, GCTA-*α*** |
| **Other European** | - | - | - | 109.138 | 111.692 | 115.714 | 118.9 | 120.646 | 121.098 | 120.58 | **-0.125, GCTA-*α*** |
| **Asian** | 22.462 | 17.332 | 9.696 | 2.472 | -2.782 | -6.09 | -8.024 | -9.074 | -9.654 | -9.744 | **-0.625, GCTA-*α*** |
| **African** | -7.622 | -9.742 | -9.216 | -13.566 | -14.638 | -14.032 | -13.464 | -13.304 | -13.422 | -13.69 | **-0.625, LDAK-thin-*α*** |
| **Mixed ancestry** | -2.506 | -24.262 | -43.002 | -50.298 | -48.648 | -43.736 | -38.786 | -34.8 | -31.842 | -30.13 | **-0.75, LDAK-thin-*α*** |

Positive value (blue) indicates GCTA**-*α*** is better over LDAK-thin**-*α*** and negative value indicates LDAK-thin**-*α*** is better.

**Supplementary Table 9: Estimated heritability (*h^2^*) and cross-ancestry genetic correlations (*r_g_*) from 4 existing methods when simulation and estimation models agree (*α*= -0.5 for all the ancestry group).**

|  |  | **Bivariate simulation combining white British and African ancestry cohort** | | | **Bivariate simulation combining white British and Asian ancestry cohort** | | | **Bivariate simulation combining Asian and African ancestry cohort** | | |
| --- | --- | --- | --- | --- | --- | --- | --- | --- | --- | --- |
| **True *r_g_*** | **GRM** | **White British (*h^2^*)** | **African (*h^2^*)** | **Genetic correlation (*r_g_*)** | **White British (*h^2^*)** | **Asian (*h^2^*)** | **Genetic correlation (*r_g_*)** | **Asian (*h^2^*)** | **African (*h^2^*)** | **Genetic correlation (*r_g_*)** |
| ***r_g_* = 0** | GRM1 | 0.547±0.011 | 0.543±0.011 | -0.001±0.014 | 0.535±0.012 | 0.512±0.012 | 0.012±0.017 | 0.531±0/011 | 0.527±0.011 | -0.002±0.018 |
|  | GRM2 | 0.471±0.010 | 0.502±0.011 | -0.004±0.028 | 0.512±0.011 | 0.492±0.012 | -0.024±0.028 | 0.479±0.010 | 0.489±0.01 | 0.020±0.029 |
|  | GRM3 | 0.537±0.013 | 0.497±0.012 | 0.002±0.029 | 0.514±0.013 | 0.495±0.013 | -0.040±0.031 | 0.515±0.012 | 0.484±0.012 | 0.021±0.038 |
|  | GRM4 | 0.494±0.011 | 0.482±0.011 | -0.005±0.033 | 0.513±0.012 | 0.492±0.012 | -0.004±0.045 | 0.491±0.011 | 0.477±0.012 | 0.020±0.041 |
| ***r_g_* = 0.25** | GRM1 | 0.531±0.010 | 0.540±0.010 | 0.152±0.013 | 0.528±0.012 | 0.485±0.011 | 0.171±0.019 | 0.531±0.010 | 0.528±0.011 | 0.162±0.015 |
|  | GRM2 | 0.471±0.010 | 0.502±0.011 | 0.250±0.027 | 0.491±0.011 | 0.489±0.011 | 0.235±0.025 | 0.480±0.010 | 0.506±0.012 | 0.251±0.026 |
|  | GRM3 | 0.529±0.013 | 0.496±0.012 | 0.246±0.049 | 0.513±0.013 | 0.482±0.012 | 0.239±0.049 | 0.527±0.012 | 0.487±0.012 | 0.252±0.060 |
|  | GRM4 | 0.492±0.011 | 0.482±0.011 | 0.273±0.047 | 0.493±0.012 | 0.480±0.012 | 0.265±0.038 | 0.495±0.011 | 0.484±0.011 | 0.255±0.039 |
| ***r_g_* = 0.50** | GRM1 | 0.525±0.009 | 0.539±0.010 | 0.251±0.011 | 0.529±0.011 | 0.502±0.011 | 0.299±0.011 | 0.518±0.010 | 0.535±0.010 | 0.284±0.013 |
|  | GRM2 | 0.469±0.010 | 0.515±0.011 | 0.454±0.023 | 0.504±0.011 | 0.485±0.011 | 0.431±0.026 | 0.475±0.010 | 0.515±0.011 | 0.470±0.025 |
|  | GRM3 | 0.527±0.013 | 0.512±0.013 | 0.464±0.051 | 0.525±0.013 | 0.484±0.013 | 0.456±0.052 | 0.494±0.013 | 0.489±0.012 | 0.472±0.036 |
|  | GRM4 | 0.486±0.011 | 0.488±0.011 | 0.498±0.036 | 0.502±0.011 | 0.482±0.012 | 0.477±0.048 | 0.478±0.011 | 0.488±0.011 | 0.511±0.046 |
| ***r_g_* = 0.75** | GRM1 | 0.519±0.008 | 0.533±0.008 | 0.325±0.010 | 0.524±0.010 | 0.508±0.009 | 0.419±0.010 | 0.530±0.010 | 0.538±0.009 | 0.371±0.010 |
|  | GRM2 | 0.473±0.009 | 0.518±0.010 | 0.661±0.019 | 0.511±0.011 | 0.496±0.011 | 0.670±0.019 | 0.481±0.010 | 0.517±0.011 | 0.661±0.020 |
|  | GRM3 | 0.511±0.013 | 0.511±0.012 | 0.712±0.051 | 0.526±0.014 | 0.492±0.012 | 0.726±0.042 | 0.516±0.013 | 0.496±0.013 | 0.728±0.045 |
|  | GRM4 | 0.486±0.011 | 0.483±0.011 | 0.764±0.049 | 0.507±0.012 | 0.489±0.011 | 0.746±0.055 | 0.485±0.012 | 0.482±0.012 | 0.765±0.049 |
| ***r_g_* = 1** | GRM1 | 0.535±0.007 | 0.529±0.006 | 0.375±0.010 | 0.524±0.008 | 0.503±0.007 | 0.441±0.008 | 0.511±0.007 | 0.527±0.008 | 0.428±0.011 |
|  | GRM2 | 0.492±0.010 | 0.518±0.010 | 0.851±0.021 | 0.511±0.011 | 0.494±0.010 | 0.891±0.015 | 0.477±0.009 | 0.517±0.010 | 0.810±0.013 |
|  | GRM3 | 0.530±0.012 | 0.512±0.014 | 0.990±0.065 | 0.512±0.013 | 0.481±0.012 | 0.980±0.049 | 0.497±0.013 | 0.512±0.013 | 0.968±0.070 |
|  | GRM4 | 0.503±0.011 | 0.508±0.010 | 0.978±0.049 | 0.498±0.012 | 0.479±0.012 | 1.020±0.052 | 0.488±0.011 | 0.481±0.011 | 0.979±0.042 |

In our simulations, we used three combinations for estimating cross-ancestry genetic correlation (white British vs. Asian, white British vs. African and Asian vs. African ancestry cohorts). In each ancestry group, we used 500,000 SNPs that were randomly selected from HapMap3 SNPs after QC. All the heritability and genetic correlation estimations were based on 500 replicates. To simulate phenotypes, we selected a random set of 1,000 SNPs as causal variants, which were presented for both ancestry groups. We used *α* = -0.5 when scaling the causal effects by ancestry-specific allele frequency in each ancestry group. Various values of genetic correlation were considered (0, 0.25, 0.50, 0.75 and 1.0). In the estimation, the four methods (GRM1 – 4) used *α* = -0.5 (standard scale factor in GRM estimation). GRM1 and 3 used all available SNPs from both ancestry groups (791581, 812332 and 777894, respectively for each combination) whereas GRM2 and 4 used only the set of SNPs common between two ancestry groups (208419, 187668 and 222106 for a, b, and c). When scaled with *α* = -0.5, GRM1 and 2 used allele frequency averaged between two ancestry groups whereas GRM3 and 4 used ancestry-specific allele frequency estimated from each ancestry group. Values highlighted in red indicate biased estimation at 5% level of significance.

**Supplementary Table 10: Estimates of heritability (*h^2^*) and cross-ancestry genetic correlations (*r_g_*) from 4 existing methods when varying *α* values across ancestry groups.**

|  |  | **Bivariate simulation combining white British and African ancestry cohort** | | | **Bivariate simulation combining white British and Asian ancestry cohort** | | | **Bivariate simulation combining Asian and African ancestry cohort** | | |
| --- | --- | --- | --- | --- | --- | --- | --- | --- | --- | --- |
| **True *r_g_*** | **GRM** | **White British (*h^2^*)** | **African (*h^2^*)** | **Genetic correlation (*r_g_*)** | **White British (*h^2^*)** | **Asian (*h^2^*)** | **Genetic correlation (*r_g_*)** | **Asian (*h^2^*)** | **African (*h^2^*)** | **Genetic correlation (*r_g_*)** |
| ***r_g_* = 0** | GRM1 | 0.568±0.011 | 0.645±0.012 | 0.0002±0.013 | 0.559±0.013 | 0.479±0.011 | 0.026±0.017 | 0.510±0.011 | 0.615±0.011 | 0.0003±0.016 |
|  | GRM2 | 0.496±0.010 | 0.504±0.011 | 0.001±0.030 | 0.516±0.012 | 0.457±0.011 | 0.033±0.031 | 0.459±0.010 | 0.501±0.011 | -0.013±0.028 |
|  | GRM3 | 0.552±0.012 | 0.671±0.011 | 0.004±0.022 | 0.538±0.013 | 0.459±0.012 | 0.035±0.039 | 0.487±0.012 | 0.636±0.012 | -0.004±0.041 |
|  | GRM4 | 0.513±0.011 | 0.636±0.011 | 0.001±0.028 | 0.518±0.012 | 0.455±0.012 | 0.030±0.035 | 0.463±0.011 | 0.603±0.011 | -0.002±0.053 |
| ***r_g_* = 0.25** | GRM1 | 0.584±0.012 | 0.647±0.011 | 0.133±0.012 | 0.569±0.012 | 0.482±0.011 | 0.164±0.016 | 0.505±0.011 | 0.617±0.011 | 0.200±0.025 |
|  | GRM2 | 0.512±0.010 | 0.503±0.010 | 0.228±0.024 | 0.533±0.012 | 0.456±0.010 | 0.241±0.027 | 0.465±0.010 | 0.507±0.012 | 0.229±0.025 |
|  | GRM3 | 0.567±0.013 | 0.670±0.011 | 0.185±0.028 | 0.551±0.013 | 0.466±0.012 | 0.248±0.048 | 0.512±0.012 | 0.627±0.013 | 0.206±0.026 |
|  | GRM4 | 0.526±0.011 | 0.631±0.010 | 0.210±0.030 | 0.529±0.012 | 0.459±0.010 | 0.251±0.036 | 0.484±0.011 | 0.600±0.012 | 0.215±0.031 |
| ***r_g_* = 0.50** | GRM1 | 0.574±0.011 | 0.621±0.011 | 0.193±0.011 | 0.551±0.011 | 0.487±0.010 | 0.285±0.012 | 0.507±0.010 | 0.607±0.011 | 0.239±0.013 |
|  | GRM2 | 0.507±0.011 | 0.516±0.011 | 0.364±0.024 | 0.534±0.011 | 0.463±0.011 | 0.449±0.034 | 0.446±0.009 | 0.510±0.011 | 0.389±0.041 |
|  | GRM3 | 0.565±0.013 | 0.657±0.011 | 0.345±0.033 | 0.541±0.013 | 0.459±0.013 | 0.464±0.051 | 0.498±0.012 | 0.633±0.011 | 0.378±0.039 |
|  | GRM4 | 0.513±0.012 | 0.628±0.010 | 0.374±0.040 | 0.529±0.012 | 0.454±0.012 | 0.491±0.051 | 0.465±0.010 | 0.601±0.011 | 0.401±0.029 |
| ***r_g_* = 0.75** | GRM1 | 0.556±0.009 | 0.581±0.010 | 0.271±0.009 | 0.546±0.010 | 0.487±0.009 | 0.381±0.009 | 0.496±0.009 | 0.595±0.010 | 0.357±0.010 |
|  | GRM2 | 0.508±0.010 | 0.535±0.010 | 0.551±0.019 | 0.521±0.011 | 0.463±0.011 | 0.641±0.022 | 0.442±0.009 | 0.512±0.010 | 0.611±0.019 |
|  | GRM3 | 0.547±0.012 | 0.687±0.011 | 0.543±0.031 | 0.544±0.013 | 0.453±0.013 | 0.731±0.056 | 0.468±0.012 | 0.633±0.011 | 0.622±0.036 |
|  | GRM4 | 0.513±0.011 | 0.656±0.010 | 0.582±0.036 | 0.512±0.012 | 0.454±0.013 | 0.751±0.039 | 0.446±0.011 | 0.605±0.011 | 0.684±0.042 |
| ***r_g_* = 1** | GRM1 | 0.554±0.008 | 0.585±0.008 | 0.325±0.010 | 0.538±0.008 | 0.486±0.007 | 0.436±0.010 | 0.506±0.009 | 0.585±0.009 | 0.480±0.012 |
|  | GRM2 | 0.523±0.010 | 0.513±0.010 | 0.635±0.017 | 0.536±0.010 | 0.465±0.010 | 0.796±0.019 | 0.465±0.009 | 0.508±0.010 | 0.709±0.017 |
|  | GRM3 | 0.567±0.013 | 0.656±0.011 | 0.701±0.034 | 0.540±0.013 | 0.451±0.013 | 0.961±0.067 | 0.497±0.012 | 0.630±0.011 | 0.740±0.048 |
|  | GRM4 | 0.531±0.011 | 0.627±0.011 | 0.781±0.055 | 0.521±0.012 | 0.447±0.012 | 1.003±0.054 | 0.461±0.011 | 0.604±0.011 | 0.801±0.037 |

In our simulation, we used three combinations for estimating cross-ancestry genetic correlation (white British vs. Asian, white British vs. African and Asian vs. African ancestry cohorts). In each ancestry group, we used 500,000 SNPs that were randomly selected from HapMap3 SNPs after QC. All the heritability and genetic correlation estimations were based on 500 replicates. To simulate phenotypes, we selected a random set of 1,000 SNPs as causal variants, which were presented for both ancestry groups. We used various *α* values that were specific to ancestries (*α* = -0.25, -0.625 and -0.75 for white British, Asian and African ancestry cohorts, respectively) when scaling the causal effects by ancestry-specific allele frequency in each ancestry group. Various values of genetic correlation were considered (0, 0.25, 0.50, 0.75 and 1.0). In the estimation, we used existing methods (GRM1 – 4) that used the standard scale factor *α* = -0.5 in GRM estimation. GRM1 and 3 used all available SNPs from both ancestry groups (791581, 812332 and 777894, respectively for each combination) whereas GRM2 and 4 used only the set of SNPs common between two ancestry groups (208419, 187668 and 222106 for a, b, and c). When scaled with *α* = -0.5, GRM1 and 2 used allele frequency averaged between two ancestry groups whereas GRM3 and 4 used ancestry-specific allele frequency estimated from each ancestry group. Values highlighted in red indicate biased estimation at 5% level of significance.

**Supplementary Table 11**: **Estimated heritability (*h^2^*) and cross-ancestry genetic correlations (*r_g_*) from the proposed method when varying *α* values across ancestry groups.**

|  | **Bivariate simulation combining white British and African ancestry cohort** | | |
| --- | --- | --- | --- |
| **True *r_g_*** | **h^2^ (British)** | **h^2^ (African)** | **Estimated cross-ancestry *r_g_*** |
| *r_g_* = 0 | 0.499±0.011 | 0.497±0.008 | 0.005±0.029 |
| *r_g_* = 0.25 | 0.486±0.011 | 0.497±0.008 | 0.267±0.031 |
| *r_g_* = 0.50 | 0.494±0.011 | 0.495±0.009 | 0.489±0.034 |
| *r_g_* = 0.75 | 0.498±0.010 | 0.493±0.009 | 0.791±0.035 |
| *r_g_* = 1.0 | 0.501±0.011 | 0.508±0.008 | 1.061± 0.044 |
|  | **Bivariate simulation combining white British and Asian ancestry cohort** | | |
| **True *r_g_*** | **h^2^ (British)** | **h^2^ (Asian)** | **Estimated cross-ancestry *r_g_*** |
| *r_g_* = 0 | 0.501±0.012 | 0.486±0.013 | 0.017± 0.042 |
| *r_g_* = 0.25 | 0.498±0.011 | 0.485±0.013 | 0.261±0.030 |
| *r_g_* = 0.50 | 0.496±0.011 | 0.485±0.013 | 0.520± 0.044 |
| *r_g_* = 0.75 | 0.482±0.011 | 0.489±0.013 | 0.764±0.047 |
| *r_g_* = 1.0 | 0.489±0.012 | 0.511±0.012 | 1.050± 0.05 |
|  | **Bivariate simulation combining Asian and African ancestry cohort** | | |
| **True *r_g_*** | **h^2^ (Asian)** | **h^2^ (African)** | **Estimated cross-ancestry *r_g_*** |
| *r_g_* = 0 | 0.497±0.012 | 0.480±0.013 | 0.021± 0.039 |
| *r_g_* = 0.25 | 0.513±0.012 | 0.489±0.012 | 0.281± 0.040 |
| *r_g_* = 0.50 | 0.484±0.012 | 0.505±0.012 | 0.474± 0.048 |
| *r_g_* = 0.75 | 0.488±0.012 | 0.483±0.012 | 0.776±0.046 |
| *r_g_* = 1.0 | 0.505±0.012 | 0.487±0.013 | 0.982± 0.051 |

In our simulation, we used three combinations for estimating cross-ancestry genetic correlation (white British vs. Asian, white British vs. African and Asian vs. African ancestry cohorts). In each ancestry group, we used 500,000 SNPs that were randomly selected from HapMap3 SNPs after QC. To simulate phenotypes, we selected a random set of 1,000 SNPs as causal variants, which were presented for both ancestry groups. We used various *α* values that were specific to ancestries (*α* = -0.25, -0.625 and -0.75 for white British, Asian and African ancestry cohorts, respectively) when scaling the causal effects by ancestry-specific allele frequency in each ancestry group. Various values of genetic correlation were considered (0, 0.25, 0.50, 0.75 and 1.0). IN the estimation, we applied the proposed method that used ancestry-specific *α* value and ancestry-specific allele frequency in GRM estimation. GRM was estimated based common SNP between population (208419, 187668 and 222106, respectively for each combination) and was implemented MTG2-*software* ^1^.

**Supplementary Table 12: Estimates of heritability (*h^2^*) and cross-ancestry genetic correlations (*r_g_*) from simulated data using estimated *α* and proposed approach of GRM across ancestry groups (true *h^2^* = 0.5 and *r_g_* = 1)**

| **Bivariate simulation combining white British and African ancestry cohort** | | | |
| --- | --- | --- | --- |
| **Number of causal SNP** | **Estimated** $\boldsymbol{h}_{\boldsymbol{1}}^{\boldsymbol{2}}$ **(British)**  **(True** $\boldsymbol{h}_{\boldsymbol{1}}^{\boldsymbol{2}}$**= 0.5)** | **Estimated** $\boldsymbol{h}_{\boldsymbol{2}}^{\boldsymbol{2}}$ **(African)**  **(True** $\boldsymbol{h}_{\boldsymbol{2}}^{\boldsymbol{2}}$**= 0.5)** | **Estimated cross-ancestry *r_g_***  **(True *r_g_* = 1.0)** |
| 100 | 0.487±0.012 | 0.509±0.011 | 1.031±0.050 |
| 1000 | 0.501±0.011 | 0.508±0.008 | 1.061± 0.044 |
| 10000 | 0.489±0.012 | 0.505±0.009 | 1.021±0.041 |
| 100000 | 0.489±0.011 | 0.497±0.010 | 0.981±0.037 |
| **Bivariate simulation combining white British and Asian ancestry cohort** | | | |
| **Number of causal SNP** | $\mathbf{Estimated}\boldsymbol{h}_{\boldsymbol{1}}^{\boldsymbol{2}}$**(British)**  **(True** $\boldsymbol{h}_{\boldsymbol{1}}^{\boldsymbol{2}}$**= 0.5)** | **Estimated** $\boldsymbol{h}_{\boldsymbol{2}}^{\boldsymbol{2}}$ **(Asian)**  **(True** $\boldsymbol{h}_{\boldsymbol{2}}^{\boldsymbol{2}}$**= 0.5)** | **Estimated cross-ancestry *r_g_***  **(True *r_g_* = 1.0)** |
| 100 | 0.505±0.012 | 0.474±0.013 | 0.978±0.061 |
| 1000 | 0.489±0.012 | 0.511±0.012 | 1.050± 0.05 |
| 10000 | 0.499±0.011 | 0.503±0.012 | 1.023±0.042 |
| 100000 | 0.488±0.011 | 0.495±0.011 | 1.012±0.039 |
| **Bivariate simulation combining Asian and African ancestry cohort** | | | |
| **Number of causal SNP** | $\mathbf{Estimated}\boldsymbol{h}_{\boldsymbol{1}}^{\boldsymbol{2}}$**(Asian)**  **(True** $\boldsymbol{h}_{\boldsymbol{1}}^{\boldsymbol{2}}$**= 0.5)** | **Estimated** $\boldsymbol{h}_{\boldsymbol{2}}^{\boldsymbol{2}}$ **(African)**  **(True** $\boldsymbol{h}_{\boldsymbol{2}}^{\boldsymbol{2}}$**= 0.5)** | **Estimated cross-ancestry *r_g_***  **(True *r_g_* = 1.0)** |
| 100 | 0.487±0.011 | 0.488±0.010 | 0.984±0.058 |
| 1000 | 0.505±0.012 | 0.487±0.013 | 0.982± 0.051 |
| 10000 | 0.498±0.011 | 0.497±0.012 | 1.022±0.040 |
| 100000 | 0.485±0.012 | 0.501±0.011 | 0.991±0.038 |

Simulation was based on 100, 1000, 10000 and 100000 random common SNPs as causal and following estimated scaling factor (*α*) across ancestries (-0.25 for white British, -0.625 for Asian and -0.75 for African ancestry cohort). All the heritability and genetic correlation estimations were based on 500 replicates. For simulation true heritability was 0.50 for both ancestry in each combined population. GRM was estimated based on our proposed approach and was implemented MTG2^1^.

**Supplementary Table 13: Estimated cross-ancestry genetic correlations (SE) for BMI**

|  | **White British**  **(n=29,628)** | **Other European**  **(n=25,909)** | **Asian**  **(n=5,719)** | **African**  **(n=5,872)** | **Mixed ancestry**  **(n=11,267)** |
| --- | --- | --- | --- | --- | --- |
| **White British** |  | **1.081 (0.043)**  **P= 5.96e-02**  $h_{c}^{2} (WB)$= 0.222 (0.012)  $h_{c}^{2} (OE)$= 0.216 (0.013) | **0.869 (0.111)**  **P= 2.37e-01**  $h_{c}^{2} (WB)$=0.195 (0.012)  $h_{c}^{2} (As)$= 0.296 (0.055) | **0.672 (0.131)**  **P= 1.22e-02**  $h_{c}^{2} (WB)$= 0.177 (0.011)  $h_{c}^{2} (Af)$= 0.245 (0.051) | **0.884 (0.082)**  **P= 1.57e-01**  $h_{c}^{2} (WB)$= 0.160 (0.011)  $h_{c}^{2} (MA)$= 0.272 (0.029) |
| **Other European** |  |  | **0.909 (0.112)**  **P= 4.16e-01**  $h_{c}^{2} (OE)$= 0.199 (0.013)  $h_{c}^{2} (As)$= 0.312 (0.056) | **0.549 (0.134)**  **P= 7.63e-04**  $h_{c}^{2} (OE)$=0.178 (0.012)  $h_{c}^{2} (Af)$= 0.239 (0.051) | **0.913 (0.085)**  **P= 3.06e-01**  $h_{c}^{2} (OE)$=0.158 (0.012)  $h_{c}^{2} (MA)$=0.282 (0.029) |
| **Asian** |  |  |  | **1.015 (0.260)**  **P= 9.53-01**  $h_{c}^{2} (As)$=0.238 (0.051)  $h_{c}^{2} (Af)$= 0.276 (0.060) | **Cohort3 is the subset of cohort 6** |
| **African** |  |  |  |  | **0.699 (0.194)**  **P= 1.21e-01**  $h_{c}^{2} (Af)$= 0.187 (0.048)  $h_{c}^{2} (MA)$= 0.203 (0.026) |
| **Mixed ancestry** |  |  |  |  |  |

P is the *p*-value of cross ancestry genetic correlation and $h_{c}^{2}$ is the heritability estimation of common SNP during bi-variate GREML. WB, OE, As, Af and MA indicates white British, other European, Asian, African, and Mixed ancestry cohorts.

**Supplementary Table 14: Estimated cross-ancestry genetic correlations (SE) for standing height**

|  | **White British**  **(n=29,663)** | **Other European**  **(n=25,940)** | **Asian**  **(n=5,725)** | **African**  **(n=5,880)** | **Mixed ancestry**  **(n=11,280)** |
| --- | --- | --- | --- | --- | --- |
| **White British** |  | **1.010 (0.018)**  **P= 5.78e-01**  $h_{c}^{2} (WB)$= 0.499 (0.011)  $h_{c}^{2} (OE)$= 0.472 (0.012) | **0.904 (0.063)**  **P= 1.27e-01**  $h_{c}^{2} (WB)$=0.473 (0.012)  $h_{c}^{2} (As)$=0.458 (0.052) | **0.876 (0.118)**  **P= 2.93e-01**  $h_{c}^{2} (WB)$= 0.446 (0.012)  $h_{c}^{2} (Af)$= 0.216 (0.047) | **1.006 (0.056)**  **P= 9.14e-01**  $h_{c}^{2} (WB)$= 0.392 (0.011)  $h_{c}^{2} (MA)$= 0.318 (0.028) |
| **Other European** |  |  | **0.847 (0.062)**  **P= 1.35e-02**  $h_{c}^{2} (OE)$= 0.449 (0.0130)  $h_{c}^{2} (As)$= 0.483 (0.054) | **0.877 (0.118)**  **P= 2.97e-01**  $h_{c}^{2} (OE)$=0.426 (0.013)  $h_{c}^{2} (Af)$= 0.224 (0.047) | **0.979 (0.057)**  **P= 7.12e-01**  $h_{c}^{2} (OE)$=0.365 (0.012)  $h_{c}^{2} (MA)$= 0.334 (0.029) |
| **Asian** |  |  |  | **0.356 (0.169)**  **P= 1.38e-04**  $h_{c}^{2} (As)$=0.452 (0.053)  $h_{c}^{2} (Af)$= 0.219 (0.049) | **Cohort3 is the subset of cohort 6** |
| **African** |  |  |  |  | **0.512 (0.158)**  **P= 2.01e-03**  $h_{c}^{2} (Af)$=0.191 (0.046)  $h_{c}^{2} (MA)$=0.273 (0.027) |
| **Mixed ancestry** |  |  |  |  |  |

P is the *p*-value of cross ancestry genetic correlation when testing that $r_{g}$=1 and $h_{c}^{2}$ is the heritability estimation of common SNP during bi-variate GREML. WB, OE, As, Af and MA indicates white British, other European, Asian, African, and Mixed ancestry cohorts.

**Supplementary Table 15: Estimated cross-ancestry genetic correlations (SE) for waist circumference**

|  | **White British**  **(n=29,666)** | **Other European**  **(n=25,946)** | **Asian**  **(n=5,807)** | **African**  **(n=5,893)** | **Mixed ancestry**  **(n=11,375)** |
| --- | --- | --- | --- | --- | --- |
| **White British** |  | **1.056 (0.052)**  **P= 2.81e-01**  $h_{c}^{2} (WB)$= 0.195 (0.0116)  $h_{c}^{2} (OE)$= 0.178 (0.0126) | **0.908 (0.145)**  **P= 5.25e-01**  $h_{c}^{2} (WB)$= 0.169 (0.0113)  $h_{c}^{2} (As)$= 0.222 (0.0554) | **0.627 (0.144)**  **P=** **9.58e-03**  $h_{c}^{2} (WB)$= 0.149 (0.011)  $h_{c}^{2} (Af)$= 0.224 (0.050) | **0.936 (0.107)**  **P= 5.49e-01**  $h_{c}^{2} (WB)$=0.135 (0.010)  $h_{c}^{2} (MA)$= 0.201 (0.029) |
| **Other European** |  |  | **1.068 (0.161)**  **P=6.73e-01**  $h_{c}^{2} (OE)$=0.159 (0.0124)  $h_{c}^{2} (As)$=0.226 (0.0553) | **0.507 (0.148)**  **P= 8.65-04**  $h_{c}^{2} (OE)$=0.145 (0.012)  $h_{c}^{2} (Af)$=0.224 (0.051) | **1.036 (0.117)**  **P= 7.57e-01**  $h_{c}^{2} (OE)$= 0.124 (0.011)  $h_{c}^{2} (MA)$= 0.207 (0.029) |
| **Asian** |  |  |  | **1.299 (0.361)**  **P= 4.07e-01**  $h_{c}^{2} (As)$= 0.158 (0.050)  $h_{c}^{2} (Af)$= 0.213 (0.051) | **Cohort3 is the subset of cohort 6** |
| **African** |  |  |  |  | **0.625 (0.256)**  **P= 1.42e-01**  $h_{c}^{2} (Af)$=0.174 (0.047)  $h_{c}^{2} (MA)$= 0.136 (0.025) |
| **Mixed ancestry** |  |  |  |  |  |

P is the *p*-value of cross ancestry genetic correlation when testing that $r_{g}$=1 and $h_{c}^{2}$ is the heritability estimation of common SNP during bi-variate GREML. WB, OE, As, Af and MA indicates white British, other European, Asian, African, and Mixed ancestry cohorts.

**Supplementary Table 16: Estimated cross-ancestry genetic correlations (SE) for hip circumference**

|  | **White British**  **(n=29,677)** | **Other European**  **(n=25,946)** | **Asian**  **(n=5,807)** | **African**  **(n=5,892)** | **Mixed ancestry**  **(n=11,373)** |
| --- | --- | --- | --- | --- | --- |
| **White British** |  | **1.076 (0.046)**  **P= 9.84e-02**  $h_{c}^{2} (WB)$=0.214 (0.012)  $h_{c}^{2} (OE)$= 0.201 (0.013) | **1.099 (0.187)**  **P= 5.96e-01**  $h_{c}^{2} (WB)$=0.187 (0.011)  $h_{c}^{2} (As)$=0.184 (0.055) | **0.778 (0.141)**  **P=** **1.15e-01**  $h_{c}^{2} (WB)$= 0.169 (0.011)  $h_{c}^{2} (Af)$= 0.230 (0.045) | **0.992 (0.097)**  **P= 9.35e-01**  $h_{c}^{2} (WB)$=0.156 (0.011)  $h_{c}^{2} (MA)$= 0.222 (0.029) |
| **Other European** |  |  | **1.027 (0.168)**  **P= 8.72e-01**  $h_{c}^{2} (OE)$=0.182 (0.013)  $h_{c}^{2} (As)$= 0.204 (0.056) | **0.572 (0.143)**  **P= 2.76e-03**  $h_{c}^{2} (OE)$=0.166 (0.012)  $h_{c}^{2} (Af)$= 0.222 (0.049) | **0.956 (0.106)**  **P= 6.79e-01**  $h_{c}^{2} (OE)$=0.143 (0.012)  $h_{c}^{2} (MA)$= 0.213 (0.029) |
| **Asian** |  |  |  | **1.391 (0.396)**  **P= 3.23e-01**  $h_{c}^{2} (As)$=0.140 (0.051)  $h_{c}^{2} (Af)$= 0.217 (0.050) | **Cohort3 is the subset of cohort 6** |
| **African** |  |  |  |  | **0.721 (0.221)**  **P= 2.06e-01**  $h_{c}^{2} (Af)$=0.178 (0.047)  $h_{c}^{2} (MA)$= 0.159 (0.026) |
| **Mixed ancestry** |  |  |  |  |  |

P is the *p*-value of cross ancestry genetic correlation when testing that $r_{g}$=1 and $h_{c}^{2}$ is the heritability estimation of common SNP during bi-variate GREML. WB, OE, As, Af and MA indicates white British, other European, Asian, African, and Mixed ancestry cohorts.

**Supplementary Table 17: Estimated cross-ancestry genetic correlations (SE) for waist-hip ratio**

|  | **White British**  **(n=29,664)** | **Other European**  **(n=25,941)** | **Asian**  **(n=5,807)** | **African**  **(n=5,095)** | **Mixed ancestry**  **(n=11,373)** |
| --- | --- | --- | --- | --- | --- |
| **White British** |  | **1.049 (0.069)**  **P= 4.77e-01**  $h_{c}^{2} (WB)$= 0.152 (0.011)  $h_{c}^{2} (OE)$= 0.139 (0.012) | **0.765 (0.179)**  **P= 1.89e-01**  $h_{c}^{2} (WB)$=0.135 (0.011)  $h_{c}^{2} (As)$= 0.163 (0.057) | NA | **0.977 (0.179)**  **P= 8.97e-01**  $h_{c}^{2} (WB)$= 0.099 (0.010)  $h_{c}^{2} (MA)$= 0.111 (0.028) |
| **Other European** |  |  | **0.921 (0.206)**  **P= 7.01e-01**  $h_{c}^{2} (OE)$=0.127 (0.012)  $h_{c}^{2} (As)$= 0.164 (0.057) | NA | **0.991 (0.189)**  **P= 9.60e-01**  $h_{c}^{2} (OE)$=0.095 (0.011)  $h_{c}^{2} (MA)$= 0.112 (0.028) |
| **Asian** |  |  |  | NA | **Cohort3 is the subset of cohort 6** |
| **African** |  |  |  |  | NA |
| **Mixed ancestry** |  |  |  |  |  |

P is the *p*-value of cross ancestry genetic correlation when testing that $r_{g}$=1 and $h_{c}^{2}$ is the heritability estimation of common SNP during bi-variate GREML. WB, OE, As, Af and MA indicates white British, other European, Asian, African, and Mixed ancestry cohorts. Some of the cross-ancestry genetic correlation was estimated as NA in the pairs involving African and this is because of no significant estimation of heritability in African ancestry cohorts.

**Supplementary Table 18: Estimated cross-ancestry genetic correlations (SE) for weight**

|  | **White British**  **(n=29,632)** | **Other European**  **(n=25,919)** | **Asian**  **(n=5,802)** | **African**  **(n=5,884)** | **Mixed ancestry**  **(n=11,360)** |
| --- | --- | --- | --- | --- | --- |
| **White British** |  | **1.062 (0.039)**  **P= 1.11e-01**  $h_{c}^{2} (WB)$=0.2608 (0.0118)  $h_{c}^{2} (OE)$=0.2351 (0.0129) | **0.891 (0.099)**  **P= 2.71e-01**  $h_{c}^{2} (WB)$=0.2339 (0.0117)  $h_{c}^{2} (As)$= 0.3235 (0.0549) | **0.832 (0.137)**  **P= 2.20e-01**  $h_{c}^{2} (WB)$= 0.215 (0.011)  $h_{c}^{2} (Af)$= 0.223 (0.049) | **0.950 (0.077)**  **P= 5.16e-01**  $h_{c}^{2} (WB)$=0.1926 (0.0108)  $h_{c}^{2} (MA)$=0.2728 (0.0292) |
| **Other European** |  |  | **0.956 (0.101)**  **P= 6.63e-01**  $h_{c}^{2} (OE)$=0.2171 (0.0128)  $h_{c}^{2} (As)$=0.3486 (0.0549) | **0.624 (0.139)**  **P= 6.83e-03**  $h_{c}^{2} (OE)$=0.197 (0.012)  $h_{c}^{2} (Af)$= 0.216 (0.049) | **0.957 (0.083)**  **P= 6.04e-01**  $h_{c}^{2} (OE)$=0.1734 (0.0117)  $h_{c}^{2} (MA)$=0.2765 (0.0294) |
| **Asian** |  |  |  | **1.054 (0.254)**  **P= 8.31e-01**  $h_{c}^{2} (As)$=0.259 (0.051)  $h_{c}^{2} (Af)$=0.223 (0.051) | **Cohort3 is the subset of cohort 6** |
| **African** |  |  |  |  | **0.670 (0.194)**  **P= 8.88e-02**  $h_{c}^{2} (Af)$=0.182 (0.047)  $h_{c}^{2} (MA)$=0.204 (0.026) |
| **Mixed ancestry** |  |  |  |  |  |

P is the *p*-value of cross ancestry genetic correlation when testing that $r_{g}$=1 and $h_{c}^{2}$ is the heritability estimation of common SNP during bi-variate GREML. WB, OE, As, Af and MA indicates white British, other European, Asian, African, and Mixed ancestry cohorts.

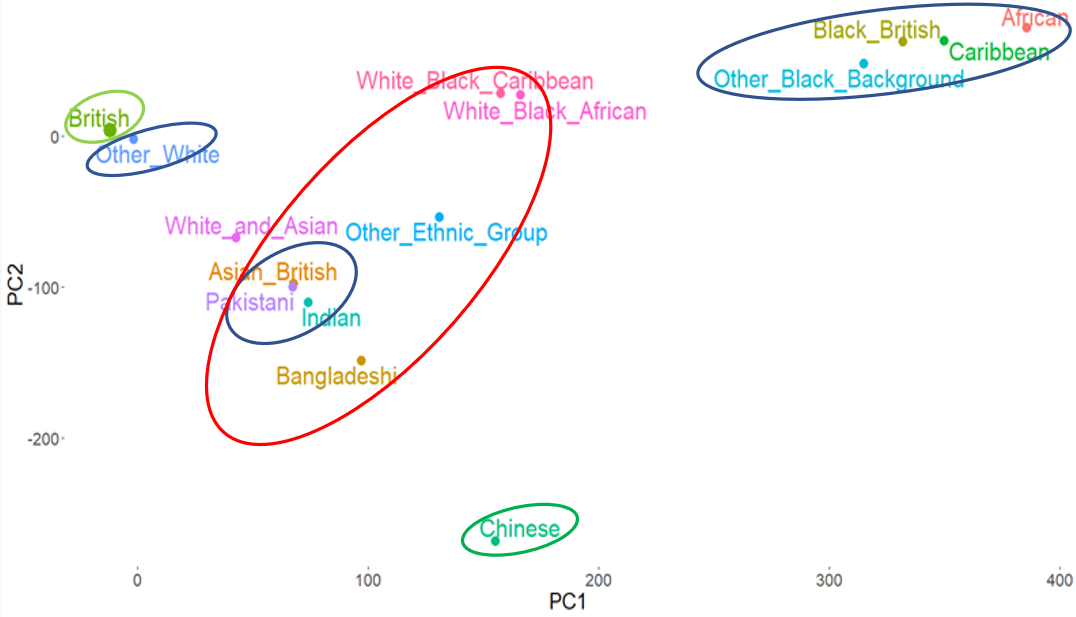

**Supplementary Figure 1: Two-dimensional scatter plots using PC1 and PC2 for UK Biobank samples.**

Individuals of the Asian ancestry are the subset of mixed ancestry cohort.

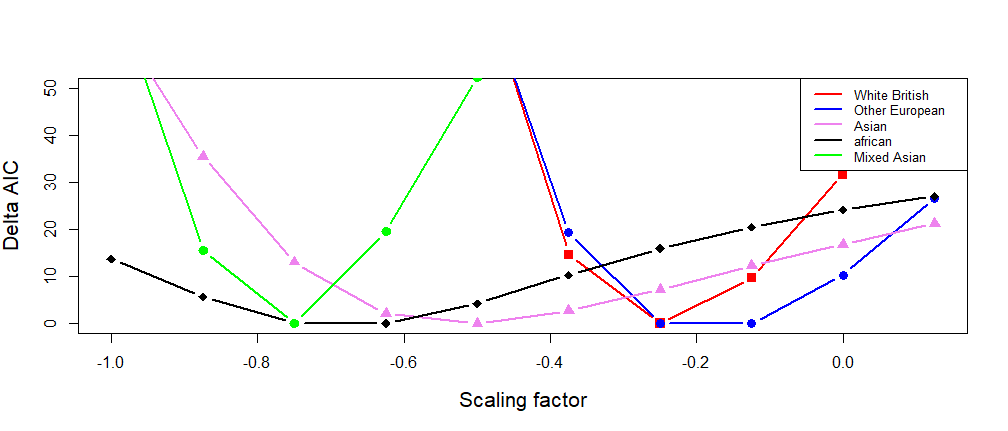

**Supplementary Figure 2: Determining optimal scaling factors (*α*) across different ancestry groups using LDAK-thin-*α* model.** LDAK-thin-*α* model assumes that SNPs contribute unequally to the heritability estimation according to their LD structure. ΔAIC values from LDAK-thin -*α* models are plotted against scaling factors, *α*, for each ancestry group. The lowest AIC (i.e. ΔAIC=0) indicates the best model. The sample sizes are 30,000, 26,457, 6,199, 6,179 and 11,797 for white British, other European, Asian, African, and mixed ancestry groups, respectively.

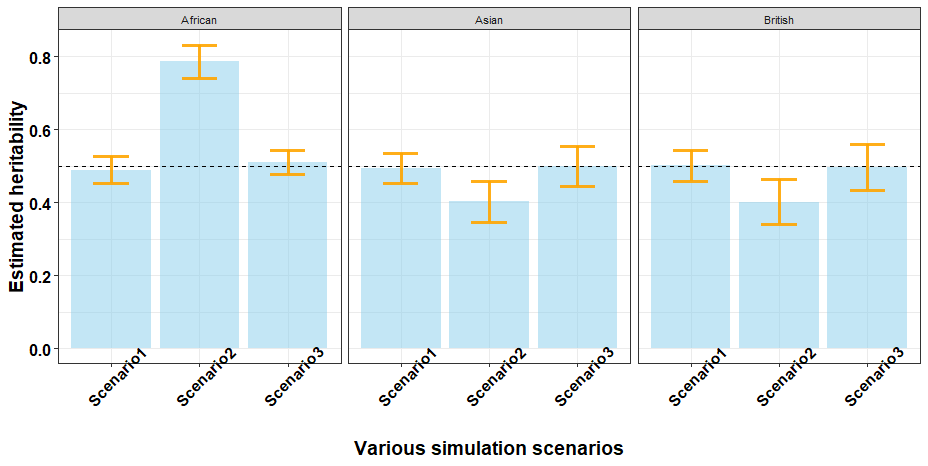

**Supplementary Figure 3:** **Estimated heritability based on simulated phenotypes for several ancestry groups.** The true heritability was 0.5 (horizontal dashed line) for simulated phenotypes using the real genotypic data after quality control in three different scenarios. Scenario 1 (simulation based	*α*= -0.5 and GRM estimated based on *α*= -0.5), scenario 2 (simulation based	*α*= -1.0 and GRM estimated based on *α*= -0.5), scenario 3 (simulation based	*α*= -1.0 and GRM estimated based on *α*= -1.0). Height of each bar is the estimated heritability and error bar indicates 95% confidence interval (CI) on 500 replicates. For simulation, the total number of individuals was 1000 and the total number of SNPs was 500,000 in each ancestry (white British, Asian and African ancestry cohorts). Simulation was based on 1000 random common SNPs as causal. The causal effect of 1000 random SNPs was estimated following a normal distribution with mean 0 for the phenotypes within ancestry group and causal effects of each SNPs were scaled by reference allele frequencies. All simulation process was implemented in MTG2. GRM for scenario1 and scenario 2 were implemented in PLINK^2^ and GRM for scenario3 was estimated in LDAK software^3^.

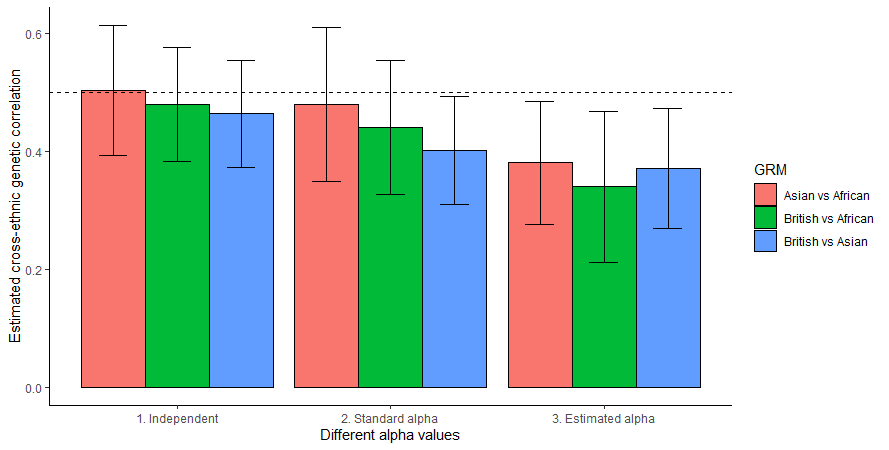

**Supplementary Figure 4: Estimated cross-ancestry genetic correlations (*r_g_*) from bivariate simulation using Popcorn across various combinations of ancestry group.** In our simulation, we used three combination for estimating cross-ancestry genetic correlation (white British with Asian, white British with African and Asian with African ancestry cohorts). Height of each bar is the estimated cross-ancestry genetic correlation and error bar indicates 95% confidence interval (CI) for 500 replicates. In the simulation, we selected a random set of 1,000 SNPs from the common SNPs as causal variant (208419, 187668 and 222106, respectively for each combination) and the true genetic correlation was 0.5. In bivariate simulation for each combination, we considered *α*= 0 for both population in independent model (first approach). In second approach, we used *α* = -0.5 (standard) for both populations, white in third approach we used the realistic *α* values (*α*= -0.25 for white British, *α*= -0.625 for Asian and *α*= -0.75 for African).

| a) Estimated cross-ancestry genetic correlations of standing height using proposed method | b) Estimated cross-ancestry genetic correlations of standing height using existing method |
| --- | --- |
| 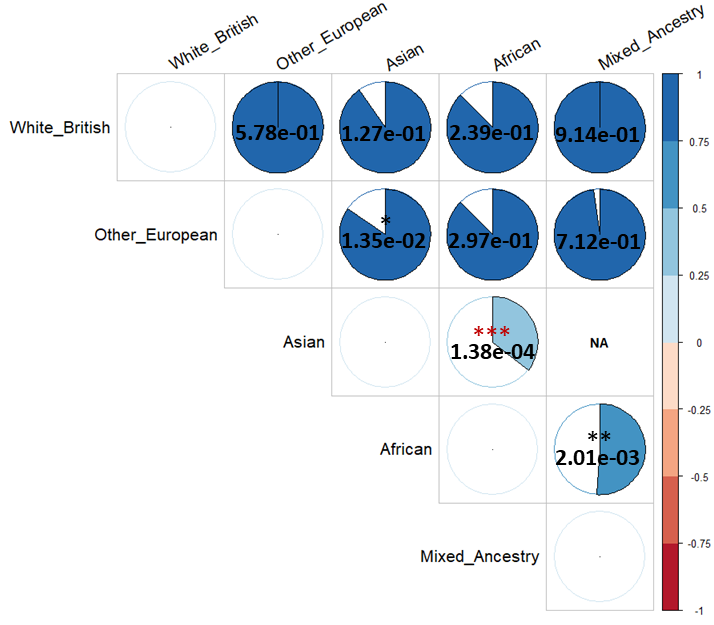 | 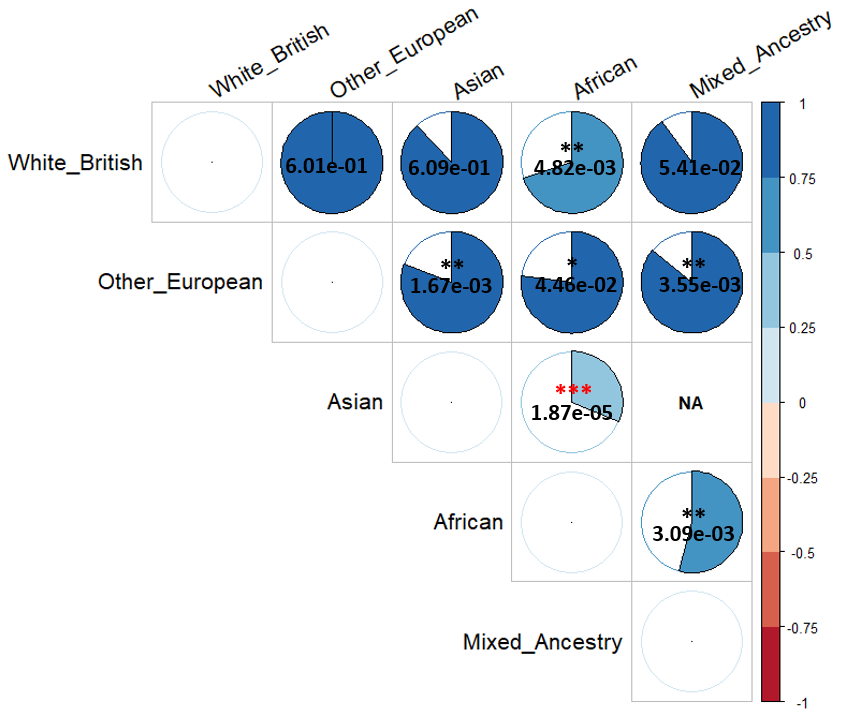 |

**Supplementary Figure 5:** Comparison of the estimations of cross-ancestry genetic correlations of standing height between proposed and existing methods. The colour and size of each pie chart indicates the magnitude of estimated cross-ancestry genetic correlations. The value in each pie chart is a *p*-value testing the null hypothesis of $r_{g}$=1. *, **, *** indicates p value < 0.05, < 0.01 and <0.001, respectively. Over-interpretation (African and European; other European and mixed ancestry) in existing method is due inaccurate consideration of *α*. Coloured asterisk indicates significantly different from 1 after Bonferroni correction (0.05/54).

| i)  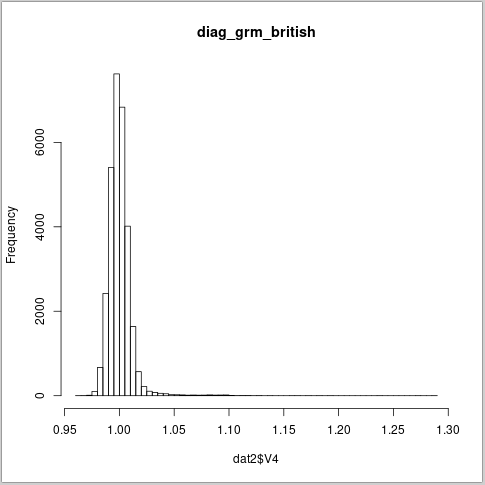 | ii)  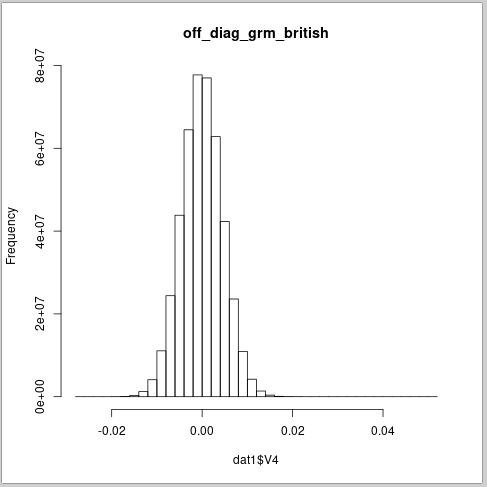 |
| --- | --- |
| **Supplementary Figure 6: Distribution of diagonal values (i) and off diagonal values (ii) of GRM (GCTA model with *α* = -0.5) in white British ancestry cohort.** Means of diagonal and off diagonal of elements are 1.0002 and -3.33e-05, respectively. | |
| i)  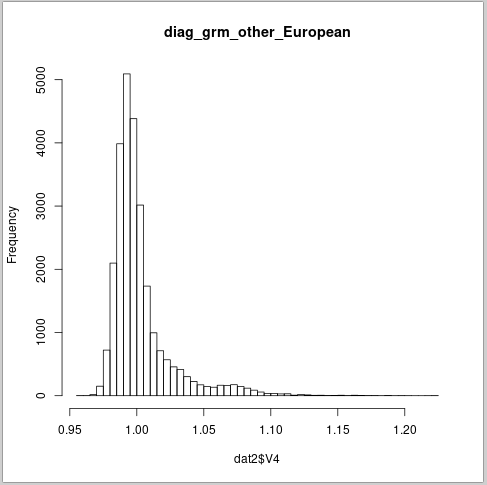 | ii)  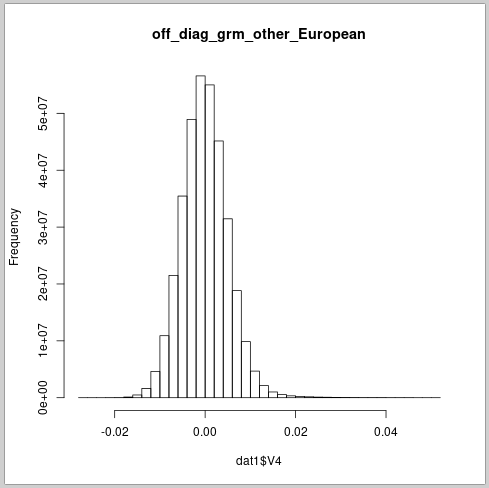 |
| i)  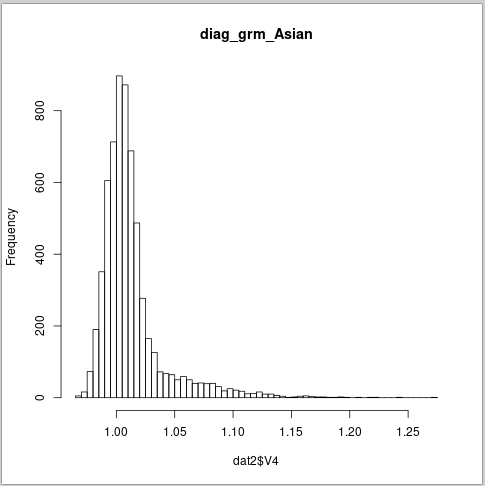 | ii)  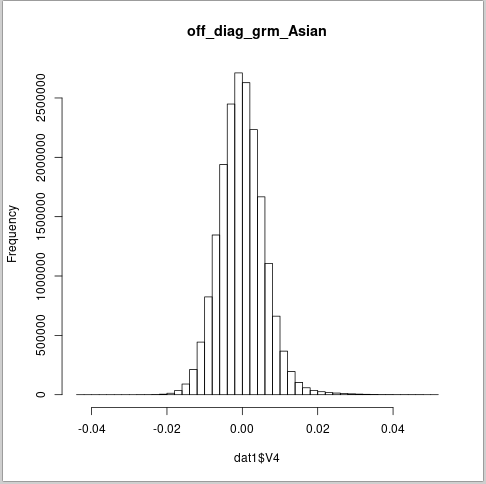 |
| **Supplementary Figure 8:** **Distribution of diagonal values (i) and off diagonal values (ii) of GRM (GCTA model with *α* = -0.5) in Asian ancestry cohort.** Means of diagonal and off diagonal of elements are 1.0128 and -1.61e-04, respectively. | |
| i)  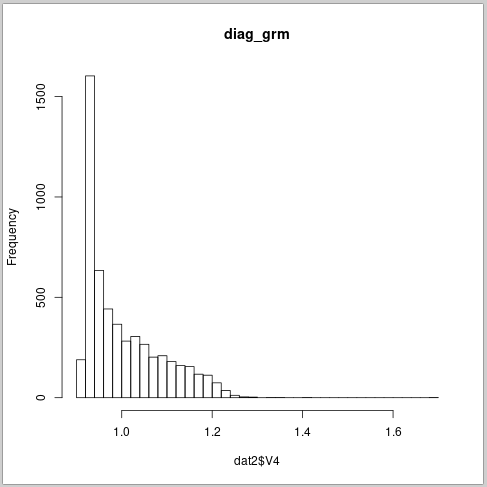 | ii)  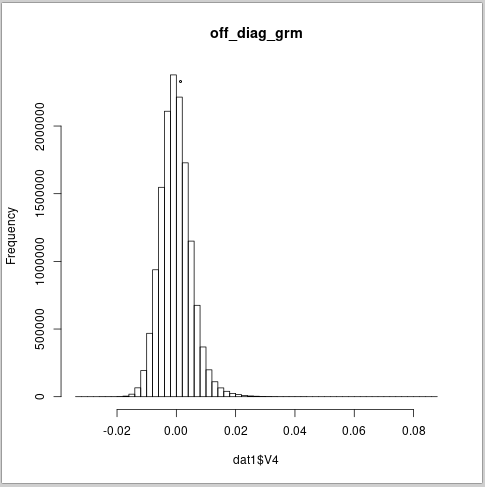 |
| **Supplementary Figure 9 Distribution of diagonal values (i) and off diagonal values (ii) of GRM (GCTA model with *α* = -0.5) in African ancestry cohort.** Means of diagonal and off diagonal of elements are 1.0033 and -1.65e-04, respectively**.** | |
| i)  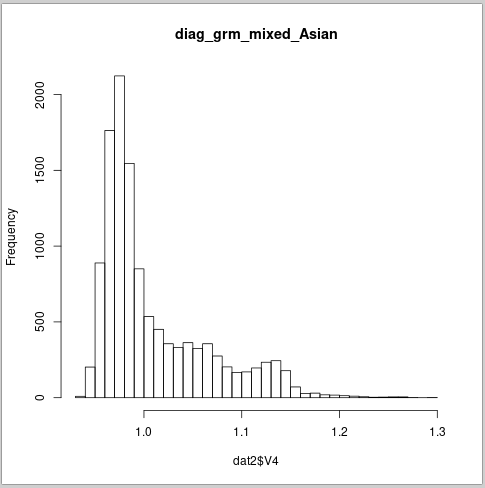 | ii)  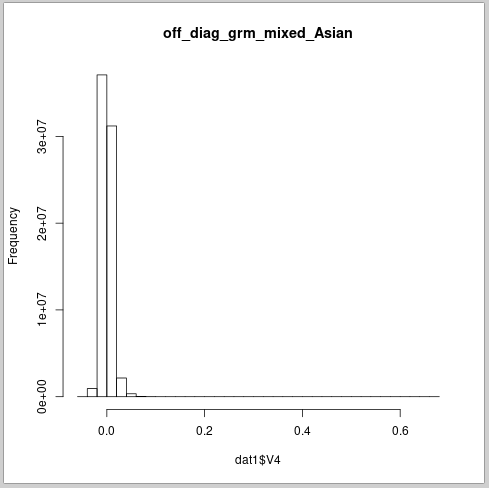 |
| **Supplementary Figure 10:** **Distribution of diagonal values (i) and off diagonal values (ii) of GRM (GCTA model with *α* = -0.5) in mixed ancestry cohort.** Means of diagonal and off diagonal of elements are 1.0096 and -8.57e-04, respectively. | |
